# Dissection and Integration of Bursty Transcriptional Dynamics for Complex Systems

**DOI:** 10.1101/2023.06.13.544828

**Authors:** Cheng Frank Gao, Suriyanarayanan Vaikuntanathan, Samantha J. Riesenfeld

**Affiliations:** Department of Chemistry, University of Chicago, IL; Institute for Biophysical Dynamics, University of Chicago, IL; Pritzker School of Molecular Engineering, University of Chicago, IL; Department of Medicine, University of Chicago, IL; Committee on Immunology, University of Chicago, IL

## Abstract

RNA velocity estimation is a potentially powerful tool to reveal the directionality of transcriptional changes in single-cell RNA-seq data, but it lacks accuracy, absent advanced metabolic labeling techniques. We developed a novel approach, *TopicVelo*, that disentangles simultaneous, yet distinct, dynamics by using a probabilistic topic model, a highly interpretable form of latent space factorization, to infer cells and genes associated with individual processes, thereby capturing cellular pluripotency or multifaceted functionality. Focusing on process-associated cells and genes enables accurate estimation of process-specific velocities via a master equation for a transcriptional burst model accounting for intrinsic stochasticity. The method obtains a global transition matrix by leveraging cell topic weights to integrate process-specific signals. In challenging systems, this method accurately recovers complex transitions and terminal states, while our novel use of first-passage time analysis provides insights into transient transitions. These results expand the limits of RNA velocity, empowering future studies of cell fate and functional responses.

## Introduction

One of the key challenges in single-cell data science, *trajectory inference* (TI) leverages genome-wide transcriptional profiles to estimate the position of each cell in an underlying, ordered biological process [1–3]. While embryonic development and cellular development are common applications, trajectory inference is also important in the analysis of other dynamic processes, such as immune responses and tumorigenesis [4–6]. The destructive nature of single-cell RNA-sequencing (scRNA-seq) technologies limits the input data to static snapshots, rather than temporal records. Computational innovations glean true dynamic information by exploiting inadvertently captured reads from unspliced pre-mRNA, as well as targeted reads from mature, spliced mRNA, to model the transcriptional kinetics of genes and thereby estimate a time derivative of the transcriptional state, known as *RNA velocity* [7, 8].

Unlike similarity-based “pseudotime” TI methods (e.g., [9–11], reviewed in [3]), RNA velocity reveals the directions and patterns of complex flows, even within a single time point, and thus also precursor and terminal cell populations. The unique capabilities and possible extensions of RNA velocity make it a potentially powerful tool in the analysis of diverse dynamic biological systems, particularly when there is limited prior knowledge. Yet, despite the advances, the effective application of RNA velocity for TI has been impeded by a lack of robustness and accuracy driven by multiple factors [12–16]. Recent approaches have used a variety of techniques to improve RNA velocity [17–26], but they do not account for distinct processes, beyond lineages, that occur simultaneously, or pluripotency. Moreover, most methods are based on ordinary differential equations and do not model intrinsic transcriptional stochasticity. The persistent gap between the promise and reality of RNA velocity has largely restricted its application.

To create an effective RNA velocity tool for investigating complex systems, such as immune responses, we created *TopicVelo*, a novel approach that disentangles potentially simultaneous processes using a probabilistic topic model [27, 28], also known as a grade-of-membership model [29, 30], which is a highly interpretable, Bayesian non-negative matrix factorization. Focusing on the specific cells and genes involved in distinct processes enables us to better capture distinct dynamics. To infer kinetic parameters for process-specific genes, *TopicVelo* fits integer transcript counts to a physically meaningful transcriptional burst model [31]. Based on the extent to which each cell participates in each process, *TopicVelo* integrates the process-specific dynamics to infer a global model of cell transitions (Fig. 1).

**Figure 1:**
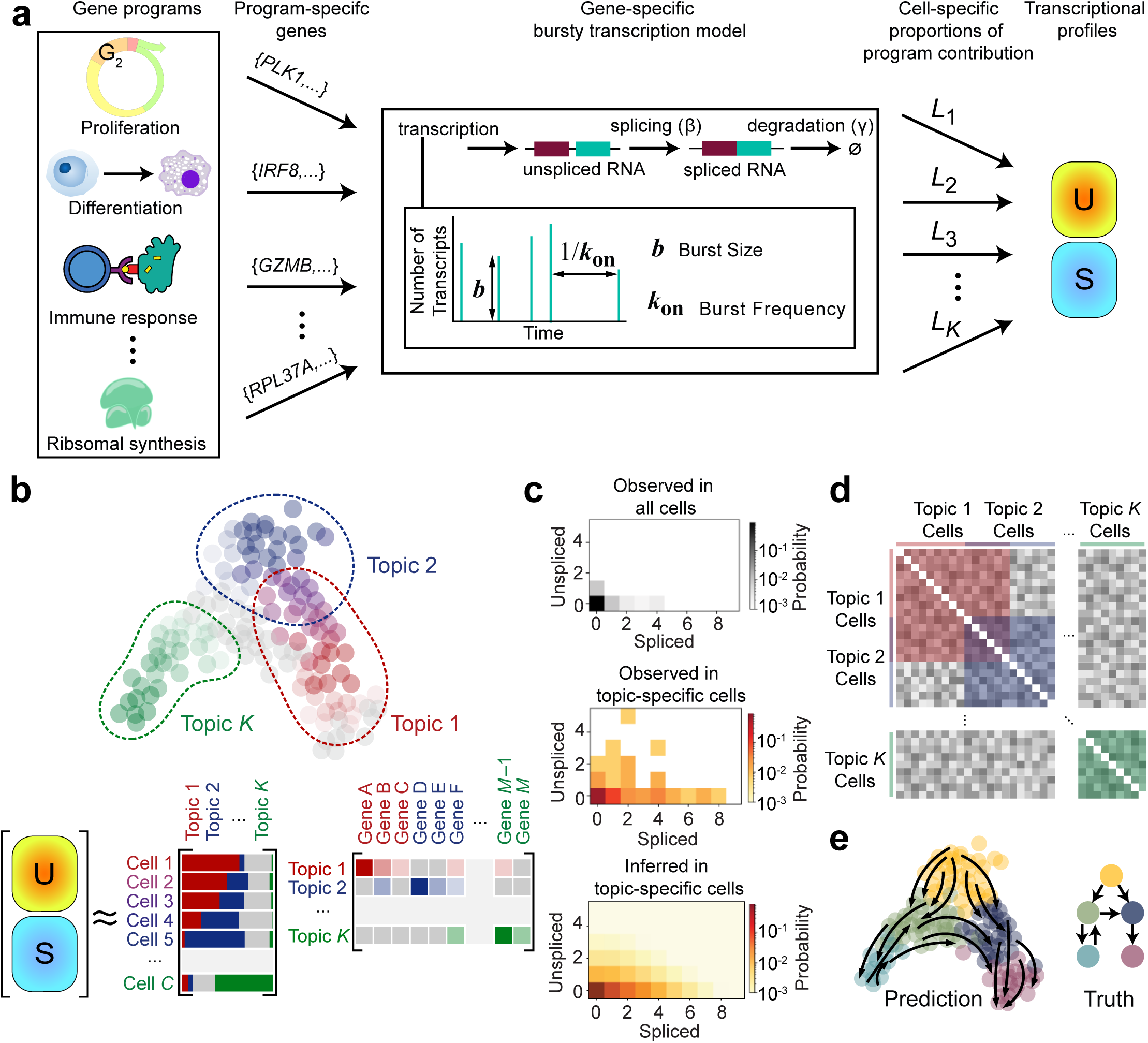
*TopicVelo* combines topic modeling and a burst model for accurate, robust RNA velocity inference. **a,** The generative model motivating *TopicVelo* accounts for distinct stochastic dynamics of transcriptional processes for different gene programs (left). Program- and gene-specific transcription follows a bursty transcriptional model governed by several parameters: the typical burst frequency *k*_on_, the burst size *b*, which has a geometric distribution, the splicing rate parameter *β*, and the degradation rate *γ* (middle). By accounting for the varying activity levels of each program *i* across cells (*L*_*i*_), the transcriptional profiles can be generated and characterized by the matrices *U* and *S*, specifying the number of unspliced and spliced transcripts, respectively, of all genes in all cells (right). **b,** A probabilistic topic model gives a Bayesian non-negative matrix factorization of the combined *U* and *S* matrix for a heterogeneous population of cells, which reveals distinct, possibly overlapping, cells and genes associated with underlying, individual programs, thereby capturing cellular pluripotency or multifaceted functionality. **c,** For many genes, the joint distribution over all cells of spliced and unspliced transcripts is concentrated at (0,0), as the gene is not involved in most cell states (top). Zooming in, the joint distribution of a topic-specific gene in topic-associated cells reveals detailed, process-specific dynamics (middle). To infer those dynamics, we fit the burst model of transcription by minimizing the KL divergence between inferred and experimentally observed joint distributions of spliced and unspliced transcripts (bottom). **d,** Cell-specific topic weights are leveraged to integrate process-specific transition signals into a global transition matrix. **e,** Results enable robust, accurate trajectory inference, as assessed by transition streamline visualizations, as well as by new mean first-passage time and terminal states analyses.

In addition to using standard visualizations of streamlines, we assessed RNA velocity results with Markovian techniques, including mean first passage time analyses that identify transient transitions not observed via traditional approaches. In diverse datasets, *TopicVelo* offers new insights and performs significantly better than state-of-the-art approach *scVelo* [8], without the aid of metabolic labeling or multiple time points, by recovering velocities, transition flows, and terminal states that are more consistent with known biology.

In the rest of the paper, we give an overview of *TopicVelo* and highlight its performance in a human hematopoiesis dataset, for which the correct dynamics were previously inferred only with the aid of metabolic labeling [18]. We also illustrate the capability of *TopicVelo* to handle complex developmental systems with stage-dependent dynamics [12, 32]. Lastly, we show *TopicVelo* infers validated, complex, convergent trajectories underlying the inflammatory responses of skin lymphocytes, using only a single time point [33].

### Overview of *TopicVelo* method

A single scRNA-seq snapshot may capture multiple biological processes, even within one cell type, including ubiquitous processes, such as proliferation and ribosomal synthesis, as well as system-specific processes, such as differentiation and immune responses (Fig. 1a). Each process involves a set of genes, or gene program, for which the process- and gene-specific kinetics are typically governed by a bursty transcription model [34]. The resulting transcriptional profiles of cells in the system also reflect the varying degrees to which different processes have been active in each cell up to the time of capture. These considerations are absent in existing RNA velocity approaches but must be accounted for in an accurate model of the generative processes of scRNA-seq data. The need to capture these key biological features motivated our approach to *TopicVelo*. Because the joint inference of all parameters in such a generative model may be computationally intractable, *TopicVelo* separates the inference of program-specific genes and cell-specific activity levels from the inference of kinetic parameters.

Specifically, *TopicVelo* operates in the following three stages:

#### 1. Process-specific inference

Inspired by previous works that effectively use probabilistic topic models to distinguish biologically relevant signals in scRNA-seq data [33, 35–37], we apply topic modeling to the combined unspliced and spliced transcript matrix (Fig. 1b) [38]. The result is a representation of each cell as a probability distribution over topics (gene programs, in the context of scRNA-seq), while each topic is a probability distribution over individual genes (Fig. 1b). Process-associated cells, i.e., cells with relatively high weights in a topic, and process-specific genes, determined using previous strategies [33, 37], serve as the input for inferring process-specific kinetic parameters. Within process-associated cells, process-specific genes can reveal important dynamic information that is hidden at the global scale and hence missed by existing methods (Fig. 1c).

The number of topics is a user-selected parameter, which, like clustering resolution, often has multiple, biologically meaningful settings. We explored several metrics developed in natural language processing (e.g., [39–41]) (Methods), and also used the literature to assess interpretability of topic-specific gene programs. Regardless, in our applications, we did not observe sensitivity of the overall results to the exact choice of topic number.

#### 2. Bursty transcription model

In contrast to the ODE-based one-state model underlying *scVelo*, *TopicVelo* efficiently fits a more faithful physical model that accounts for transcriptional bursting, adapting a previous model for studying mRNA transport [31] (Fig. 1a, c). The chemical master equation of the model for a given gene is:

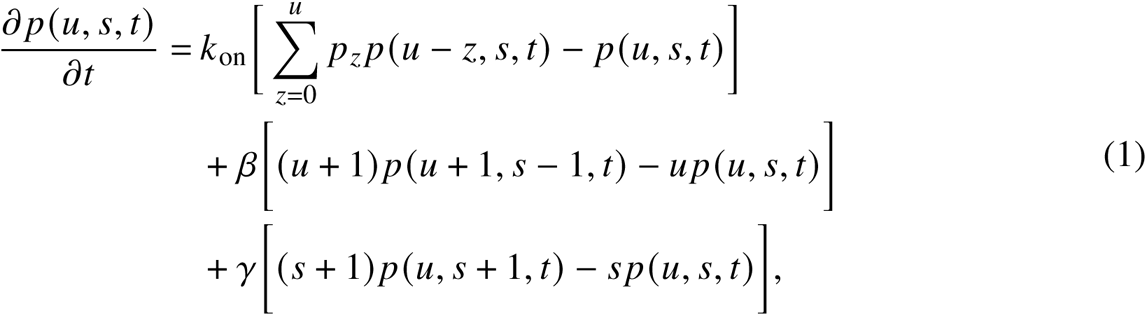

where *p* (*u*, *s*, *t*) is the probability of observing a cell with *u* unspliced pre-mRNA transcripts and *s* spliced mature mRNA transcripts at time *t*; *k*_on_ is the rate of the Poisson process governing the burst event; *p*_*z*_, the probability of producing *z* unspliced pre-mRNA transcripts during a single burst event, is governed by a geometric distribution; *β* is the splicing rate; and *γ* is the rate of degradation of spliced mRNA. Parameters are initialized with the method of moments or another heuristic. For a given parameter setting, *TopicVelo* uses an implementation of the Gillespie algorithm [42] to estimate the full joint distributions of unspliced and spliced transcript counts. Then the Nelder-Mead algorithm implemented in *SciPy* [43] is used to infer the maximum likelihood parameter values (Supplementary Fig. 1).

#### 3. Integration of process-specific dynamics

A key feature of *TopicVelo* is the capability to integrate process-specific transition matrices into a global transition matrix (Fig. 1d). First, from the inferred process-specific kinetic parameters, *TopicVelo* constructs process-specific transition matrices, based on a previous approach [8], namely by applying a exponential kernel to the cosine similarities between velocities and differences in spliced expression among nearest neighbors. Each transition matrix characterizes the probabilistic flow of process-specific transcriptional changes across process-associated cells. Then a larger scale or global transition matrix is constructed by linearly combining process-specific transition matrices, using the topic weights of cells. This strategy enables locally important dynamics to be accurately recovered and then woven into larger-scale, complex trajectories. The user-selected topic weight threshold, which determines topic-associated cells, balances an inherent trade-off between the benefit of separating dynamic processes and the risk of losing dynamic range and/or information in overlaps among topic-associated cells.

#### 4. Revealing cell state transitions by analyzing the integrated transition matrix

In addition to assessing results with typical streamline visualizations, we use the stationary distribution of the integrated transition matrix to identify terminal cell populations. Furthermore, we introduce the use of mean first passage time (MFPT) analysis to gain insights into transient transitions invisible at the global scale with traditional approaches.

We analyzed the performance of *TopicVelo* in diverse applications, detailed below, which revealed its accuracy and capacity to offer interpretable, biological insights (Fig. 1e).

### *TopicVelo* infers challenging trajectories in human hematopoiesis without metabolic labeling

RNA velocity inference without metabolic labeling is often inaccurate [18], but incorporating metabolic labeling into scRNA-seq remains an experimental challenge [44]. To test the effectiveness of *TopicVelo*, we applied it to human hematopoiesis data from a recent study in which RNA velocity was extended to leverage single-cell metabolic labeling techniques that distinguish newly synthesized versus preexisting transcripts [18]. The published analysis reconstructed a complex, multifurcating trajectory of transitions which *scVelo* fails to capture. Using *TopicVelo* on the data *without* the metabolic labels, we inferred the correct transitions, including streamlines that accurately delineate the trajectories of monocytes, basophils, erythrocytes, and megakaryocytes (Fig. 2a).

**Figure 2:**
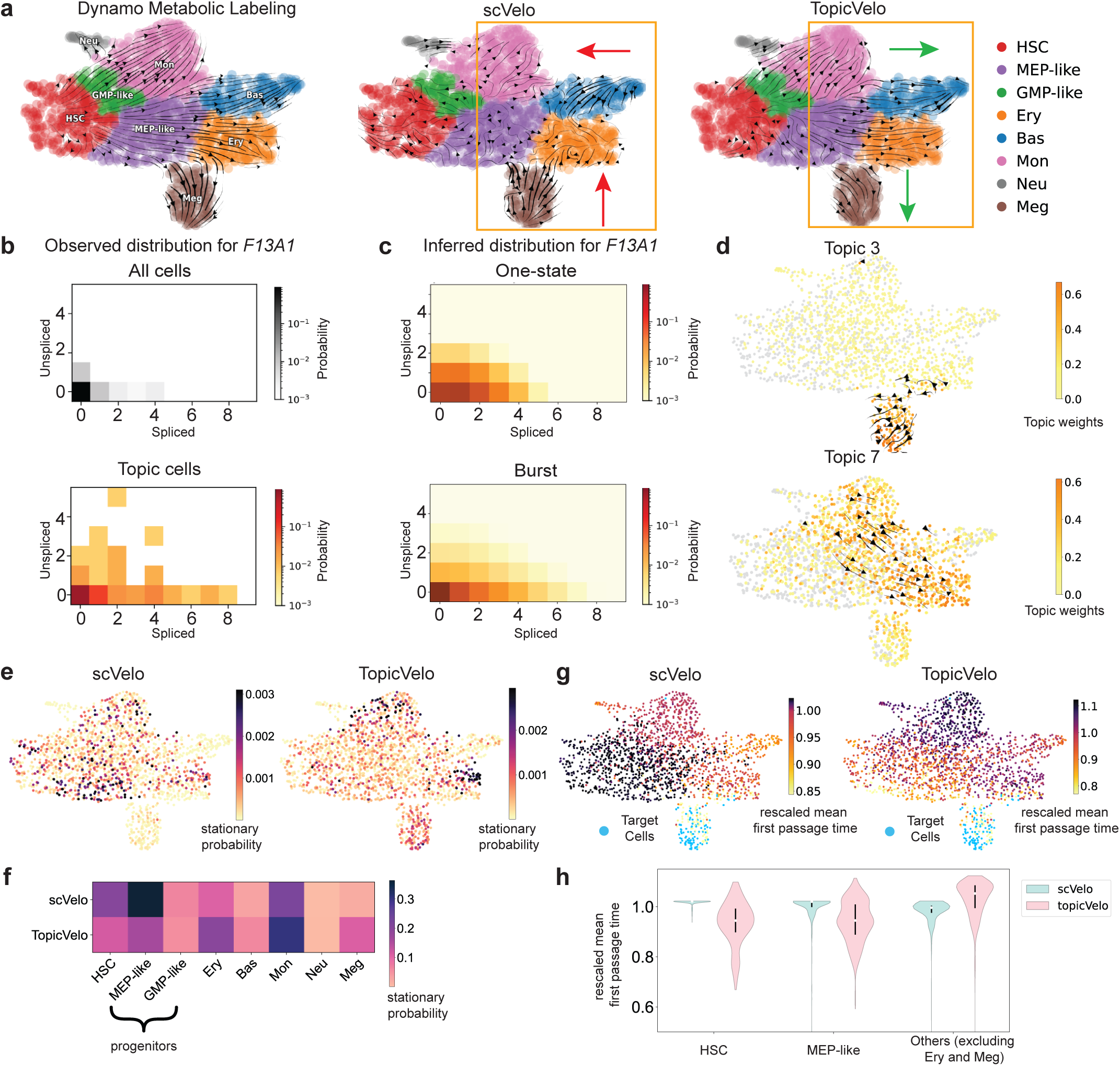
*TopicVelo* inferred multi-furcating trajectories of human hematopoiesis whose recovery previously required metabolic labeling. **a**, Previously published [18] UMAP embedding of hematopoiesis data shows cells colored by annotated progenitor (HSC, hematopoietic stem cell; MEP-like, megakaryocyte and erythrocyte progenitor; GMP-like, granulocyte and monocyte progenitor) and terminal (Ery, erythrocyte; Bas, basophil; Mon, monocyte; Neu, neutrophil; Meg, megakaryocyte) cell types. Streamlines (arrows) were inferred either with metabolic labeling, by *Dynamo* (left), or without it, by the *scVelo* dynamical model (middle), and by *TopicVelo* with an 8-topic model (right); *TopicVelo* but not *scVelo* captures key cell-type differentiation (green versus red arrows). **b,** Plots show the experimental joint distribution of spliced and unspliced mRNA counts in all cells, or cells with highest weight in topic 3, of the topic-3 specific gene *F13A1*, which is known to be expressed in megakaryocytes [45]. **c,** Plots show the joint distribution of *F13A1* in topic-3 high cells, inferred using the one-state model, or maximum likelihood estimates for the burst model; the latter better captures both the diffuseness of the joint distribution and the empirical concentration at (0,0). **d,** Topic-specific streamlines obtained from topic-specific transition matrices for topics 3 and 7 respectively. The color bar indicates the topic weights for cells used in the parameter inference. The topic 3 plot demonstrates transitions into mature megakaryocytes and the topic 7 plot suggests transitions into erythroid. e, f, *TopicVelo* identified terminal states missed by *scVelo*. UMAPs (**e**) show stationary probabilities for *scVelo* (left) and *TopicVelo* (right) transition matrices; summary heatmap (**f**) highlights relatively high probabilities from *TopicVelo* for terminal versus progenitor cell types (columns). In particular, *TopicVelo* identifies megakaryocytes as a terminal state concurring with the global and topic-specific streamlines. g, h, *TopicVelo* estimated shorter transition times for true differentiation pathways. UMAPs (**g**) show mean first-passage times to megakaryocytes (Target, blue), rescaled by median, based on *scVelo* (left) and *TopicVelo* (right); summary violin plots (**h**) highlight shorter transition times from progenitors versus others (x axis) estimated by *TopicVelo*, but not scVelo. (White dot: median, black vertical lines: 25th-75th percentile.)

To obtain global transition matrix of *TopicVelo*, we first performed topic modeling [37, 38], resulting in an 8-topic model that identifies gene programs associated with both known cell types (topics 1 and 3) and heterogeneous cell states during differentiation (Supplementary Fig. 2, Supplementary Table 1). For example, megakaryocyte-associated topic 3 appropriately features the gene *F13A1*, which codes for a subunit of plasma factor XIII known to be produced by megakaryocytes [45] (Supplementary Fig. 2d, 3a). Though a global phase plot of *F13A1* indicates little transcriptional activity, focusing on cells with highest weight in topic 3 brings the dynamical features of *F13A1* into relief (Fig. 2b).

Based on the burst model, *TopicVelo* then inferred topic-specific kinetic parameters for topic-specific genes. By assuming a steady state can be approximated by the joint distributions of spliced and unspliced counts of topic-specific genes in topic-associated cells, *TopicVelo* substantially improved upon the parameter estimates inferred from the one-state model underlying *scVelo*. For example, it more accurately recovered the experimental joint distribution of *F13A1* over topic-3 high cells (Fig. 2c). Indeed, while velocities of topic-3 specific genes *F13A1*, *PLEK*, and *ZYX* were inferred to be negative by *scVelo*, *TopicVelo* inferred them to be positive, consistent with experimental evidence that these genes are up-regulated during megakaryocytic differentiation [46, 47] (Supplementary Fig. 3a–c). Similarly, whereas *scVelo* inferred down-regulation of the basophil-associated, topic-1 specific genes *GATA2* and *HPGD*, *TopicVelo* predicted their up-regulation in the basophil lineage, consistent with previous experiments showing that *GATA2* is critical for basophil development [48] and *HPGD* is enriched in basophils [49] (Supplementary Fig. 3d, e). Using the inferred topic-specific signals, *TopicVelo* then created topic-specific transition matrices, whose corresponding streamlines were consistent with those inferred for the same regions using metabolic labeling data (Fig. 2d).

Finally, these topic-specific transition matrices were integrated to obtain the global transition matrix and corresponding streamlines (Fig. 2a). To quantitatively evaluate the quality of inference by *TopicVelo*, we computed the stationary distribution as a proxy for identifying terminal states. While both *scVelo* and *TopicVelo* assigned relatively high stationary probabilities to erythroid and monocytes, *TopicVelo* additionally recognized megakaryocytes as terminal states (Fig. 2e). Furthermore, aggregation of the stationary probabilities by cell types illustrated that, compared to *scVelo*, *TopicVelo* suggested higher stationary probability for terminal cell types and lower probability for progenitors, consistent with the expected cell-fate transitions.

To investigate the dynamics and the trajectories of differentiation, we used the MFPT to gauge the identities of ancestral populations and assess the likelihood of populations transitioning into terminal states. For instance, we computed cell-specific MFPTs to megakaryocyte-like cells and observed that the MFPTs derived from *scVelo* versus *TopicVelo* displayed very different trends (Fig. 2g). In particular, *TopicVelo* estimated lower MFPTs for progenitors than for other, non-megakaryocyte terminal cell types, whereas *scVelo* estimated the opposite. The inference from *TopicVelo* agrees better with the established biological understanding (reflected in the cell names) that megakaryocytes originate directly from progenitors, rather than from other terminally differentiated populations.

Collectively, these results demonstrate the capacity of *TopicVelo* to identify biologically meaningful dynamic genes, infer more biologically accurate RNA velocity, and provide more meaningful insights into the terminal states and trajectories of differentiation.

### *TopicVelo* recovers complex developmental trajectories in mouse and human

Several studies have observed that some genes exhibit developmental-stage dependent transcription rates, termed “multiple rate kinetics (MURK)” [8, 12–14, 32]. Moreover, *scVelo* does not account for this stage dependency and erroneously produced reversed streamlines for mouse erythropoiesis when MURK genes were included in the data [12]. In contrast, *TopicVelo* produced the correct trajectories in this setting (Fig. 3a). A stationary distribution analysis further confirmed the streamline visualization; whereas *scVelo* falsely identified intermediate erythroid stages as terminal states, *TopicVelo* correctly suggested that essentially all of the stationary probability is in the erythroid 3 cell state (Fig. 3b).

**Figure 3:**
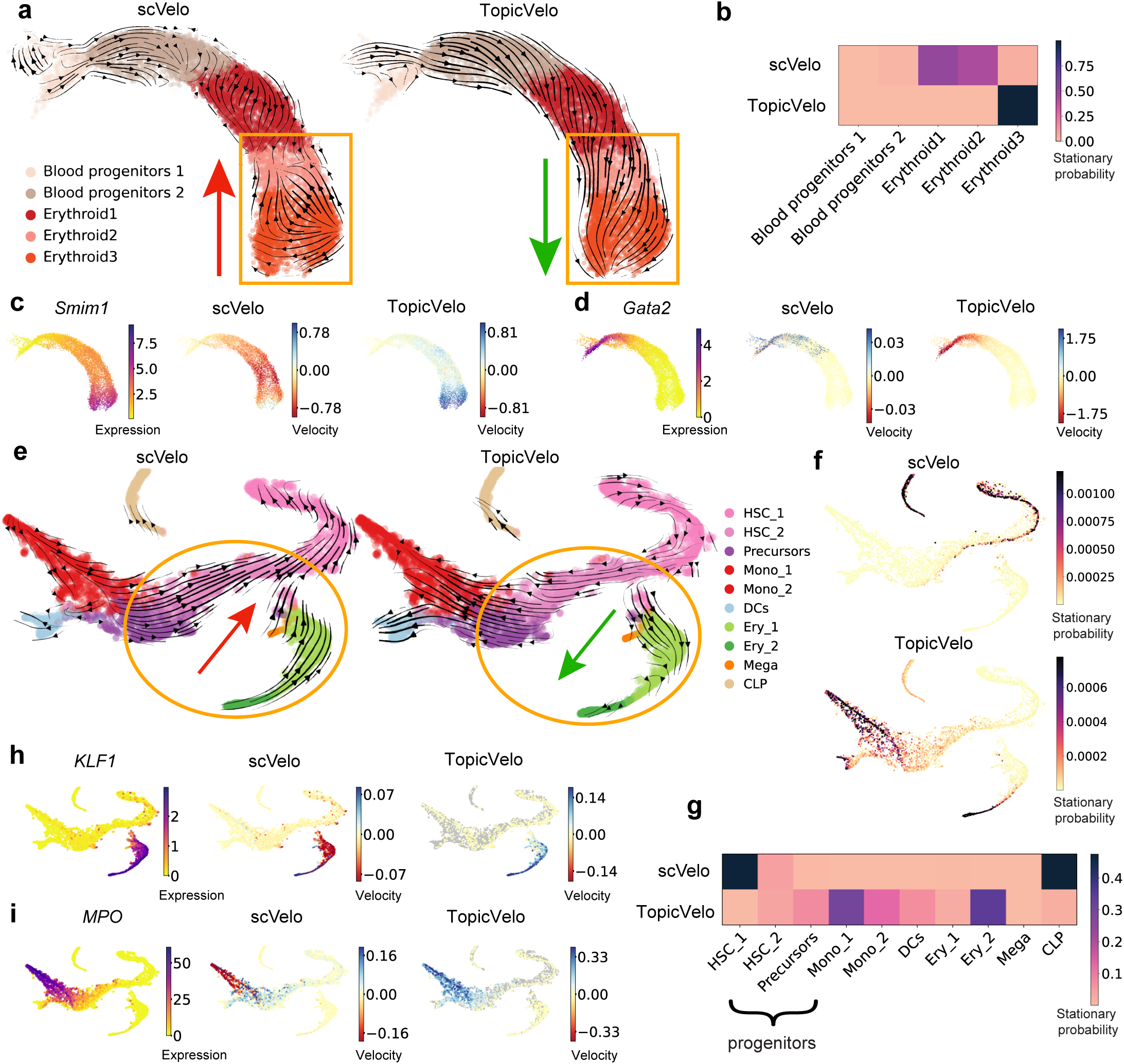
*TopicVelo* correctly captures mouse erythropoiesis and human bone marrow development trajectories. **a, b,** *TopicVelo* accurately identifies erythroid 3 as a terminal state. Previously published [12, 14] UMAP embeddings of cells in erythropoiesis, colored by cell-type annotation (**a**), shows streamlines (arrows) inferred by the *scVelo* stochastic model (left), which erroneously suggests differentiation of erythroid 3 into erythroid 2 cells (red arrow), or by *TopicVelo* (right), which recovers the expected differentiation trajectory (green arrow). Heatmap (**b**) shows the stationary probability distributions (color) from *scVelo* and *TopicVelo* (rows), aggregated by cell types (columns). **c, d,** UMAP plots for the topic-specific genes *Smim1* (**c**) and *Gata2* (**d**), with cells colored by smoothed gene expression (left), and by velocities (negative, red; positive, blue) inferred by *scVelo* (middle), or by *TopicVelo* (right). **e–g,** *TopicVelo* correctly discovers terminal cell types in human bone marrow development. Previously published [32] *t*-SNE plot of cells from human bone marrow, colored by annotated cell type (**e**), shows streamlines inferred by *scVelo* stochastic model (left), which incorrectly predicts that precursors, megakaryocytes (Mega), and erythrocytes (Ery) differentiate into hematopoietic stem cells (HSC) (red arrow), or by *TopicVelo* (right), using 10 topics, which recovers the expected trajectories for all major lineages (green arrow). (Mono: monocyte, DC: dendritic cell, CLP: common lymphoid progenitor.) Same *t*-SNE plots of cells colored by stationary probability (**f**) as inferred by *scVelo* (top) and *TopicVelo* (bottom). Heatmap (**b**) shows the stationary probability distributions from *scVelo* and *TopicVelo*, aggregated by cell types. **h, i,** *TopicVelo* gives markedly different velocity results from those of *scVelo* for topic-specific genes. For the erythroid-associated gene *KLF1* (**h**) and monocyte-associated gene *MPO* (**i**), *t*-SNE plots show cells colored by smoothed gene expression (left), and by velocities inferred by *scVelo* (middle), or by *TopicVelo* (right).

Biologically informative results were achieved by *TopicVelo* using 2 topics, which accurately modeled expression patterns during the maturation of blood progenitors to erythroid cells (Supplementary Fig. 4, Supplementary Table 1). Topic 0 has weights increasing across the developmental process and features the archetypal red blood cell genes *Hba-x* and *Hbb-y* [12], and their unspliced counterparts, as well as *Smim1*, which influences red blood cell traits [50](Fig. 3c, Supplementary Fig. 4a). Inversely, topic 1 weights decrease across the developmental process, as does the expression of topic-1 specific genes, such as *Gata2*, *Fn1* and *Fscn1* (Fig. 3d, Supplementary Fig. 4b). These results corroborate previous observations that *Gata2* is highly expressed in progenitors, with expression declining after erythroid commitment [51], and that *Ccnd2* expression is anti-correlated with erythroid progression [52].

**Figure 4:**
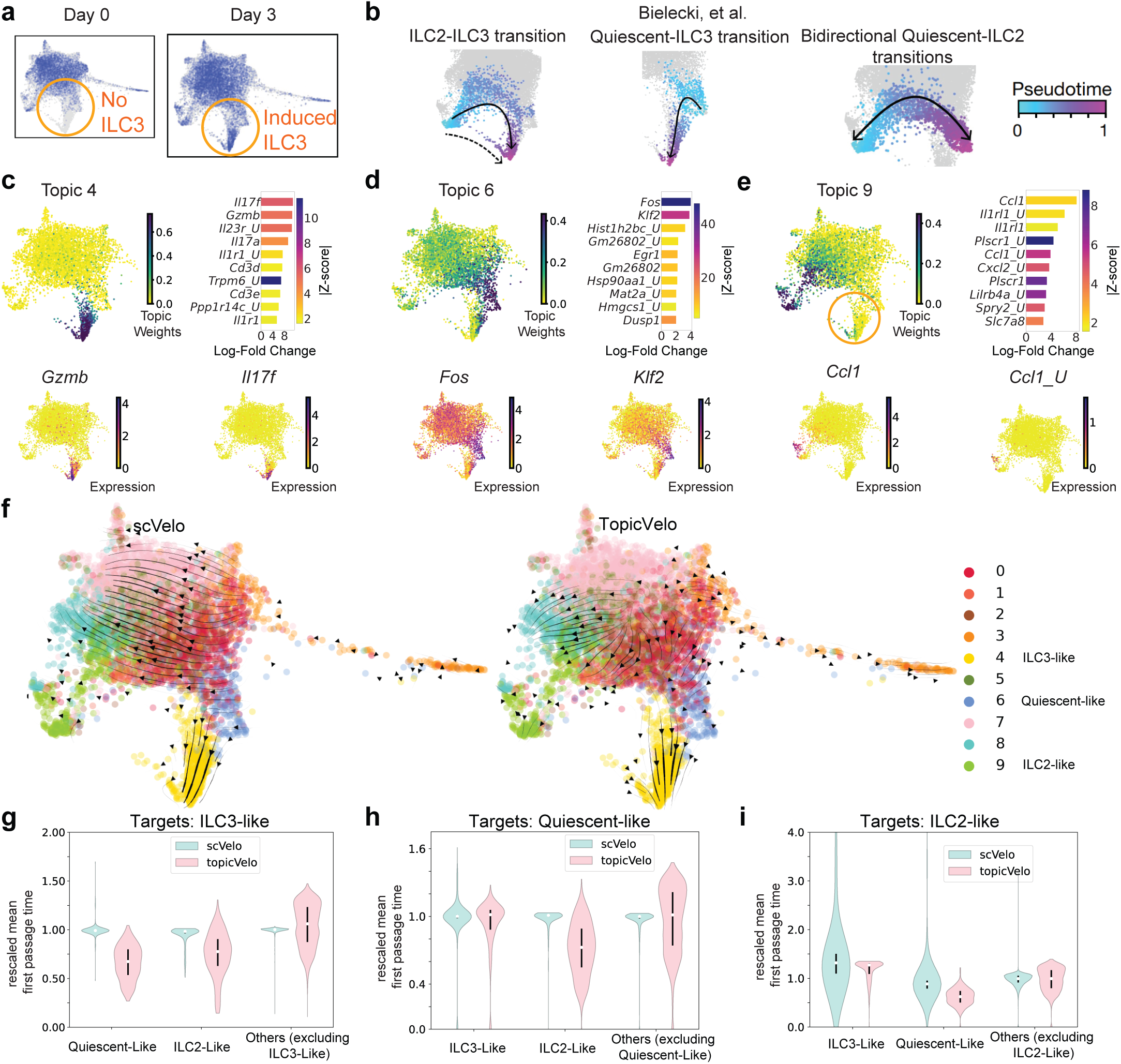
Using data from only one of five time points, *TopicVelo* reveals complex transitions underlying the inflammatory response of skin ILCs. **a, b,** Previously published [33] force-directed layout (FDL) embedding of scRNA-seq profiles of skin innate lymphoid cells (ILCs) from a mouse model of psoriasis, colored by day of collection (**a**), and by pseudotime (**b**) in independently inferred, diffusion-based trajectories (panels), with directionality (arrows) imposed by the presence of ILC3-like cells (orange circle) at day 3 but not at day 0. **c–e,** Highlights of 3 topics from a 10-topic model of both spliced and unspliced mRNA transcripts for day-3 cells, only. For ILC3-like topic 4 (**c**), quiescent-like topic 6 (**d**), and ILC2-like topic 9 (**e**), the FDL plots show day-3 cells, colored by topic weight (top left) and by the log-normalized expression of topic-specific genes (bottom left, right); bar chart (top right) shows the top 10 topic-specific genes by largest log-fold change, colored by z-score (*’ U’* appended to gene symbol indicates unspliced transcript). A subset of induced cells has relatively high topic weights for both topics 4 and 9 (orange circle, e). f–i, *TopicVelo* disentangles simultaneous but distinct dynamics of ILC responses. FDL plots of day-3 cells, colored by most strongly associated topic (**f**), show streamlines (arrows) from *scVelo* dynamical model (left) or *TopicVelo* (right), using the topic model from c–e. Focusing on the transitions to ILC3-like cells (yellow, high in topic 4, as in c), both methods predict the transition from quiescent-like cells (blue, high in topic 6, as in d), but only *TopicVelo* correctly predicts the experimentally validated transition from ILC2-like cells (green, high in topic 9, as in e) via a subset of cells high in both topics 4 and 9. Violin plots show the distributions of median-rescaled mean first passage times, estimated using *scVelo* (light blue) and *TopicVelo* (pink), from different groups of non-target cells (x axis) to different target populations, i.e., ILC3-like (**g**), quiescient-like (**h**), and ILC2-like (**i**) cells. Smaller values indicate faster inferred transition times, suggesting better support for that biological transition. (White dot: median, black vertical line: 25th–75th percentile.)

In another challenging setting involving complex, multi-furcating, human hematopoietic stem cell (HSC) differentiation [32], *TopicVelo* used a 10-topic model to recover the expected trajectories and identify key genes involved in cell-fate commitments, without the prior knowledge of starting state required by pseudotime inference (Supplementary Fig. 5) and without inferring erroneous reversals in directionality, as did *scVelo* (Fig. 3e). The stationary distribution analysis confirmed that *scVelo* incorrectly identified early stage HSCs as terminal states, whereas most of the stationary probability derived from *TopicVelo* was associated with true terminal states (Fig. 3f, g).

The inferred topics characterized different stages of development and identified key, lineage-specific genes, leading to velocity predictions that are more consistent with known biology. For example, topic 6 is relatively high in erythroid cells and includes the gene *KLF1*, previously shown to be correlated with erythroid commitment [32]. In contrast to *scVelo* predictions that early erythroid cells (Ery 1) down-regulate *KLF1*, *TopicVelo* accurately predicted that they up-regulate *KLF1* (Fig. 3h). *TopicVelo* also highlighted several other patterns previously observed in the literature, including up-regulation of *MPO* during monocyte commitment [32] (Fig. 3i), up-regulation of *CA1* in the peripheral blood erythroid cells [53], association of *IRF8* with monocyte development and dendritic cell function [54]; expression of *SELP* during megakaryocyte development [55], down-regulation of *CRHBP* in HSCs during differentiation [56], and expression of the chemotactic gene *AZU1* in monocytes [57] (Supplementary Fig. 6, Supplementary Table 1).

Together, these results indicate that *TopicVelo* outperforms the state-of-the-art in settings of complex cellular differentiation and gene expression patterns, recovering much more biologically accurate trajectories and highlighting informative genes.

### *TopicVelo* predicts bidirectional and convergent immune responses of innate lymphoid cells

An important motivation for developing *TopicVelo* was to meet the challenge of analyzing complex immune responses, including those involving unconventional trajectories, such as convergence on one cell state from multiple origins and functional plasticity between cell types [33, 58, 59]. With different gene programs involved in conversions in opposite directions between cell types, traditional approaches to RNA velocity and trajectory inference do not reveal such intricacies. We tested *TopicVelo* in this setting by analyzing our previously published scRNA-seq data from innate lymphoid cells (ILCs) isolated from the skin of mice in a model of psoriasis [33]. To disentangle transcriptional states of skin ILCs and model their trajectories, the study leverages scRNA-seq data collected from mice sacrificed at five time points (days 0–4) during the inflammatory immune response, in combination with topic modeling and density-based pseudotime inference. The detailed analysis and extensive experimental validations demonstrate multiple possible transitions to a pathogenic ILC3-like state, including an ILC2-ILC3 transition, confirmed using a transgenic fate-mapped mouse, which may occur via two routes, as well as a quiescent-ILC3 transition and possibly bidirectional quiescent-ILC2 transition (Fig. 4b).

Using data from day 3 only, we assessed the capability of *TopicVelo* to predict these complex immune response trajectories without information from multiple time points or specification of root and terminal states. Consistent with the previous analysis, a 10-topic model identified three topics strongly associated with the ILC states involved in the transitions previously analyzed (Fig. 4c–e), as well as other topics characterizing this heterogeneous ILC landscape (Supplementary Fig. 7, Supplementary Table 1). Topic 4 is strongly associated with the ILC3-like cells, and characterized by proinflammatory, ILC3- and T_H_17-associated genes, such as *Il17a*, *Il23r*, *Gzmb*, and *Il1r1* [60] (Fig. 4c). Topic 6 features a gene program previously identified as “quiescent-like” [33], including *Klf2*, a transcription factor associated with T cell quiescence [61] (Fig. 4d). Topic 9 features ILC2- and T_H_2–associated genes, such as *Il1rl1* (ST2, the receptor for IL-33) [60], as well as chemokines, such as *Ccl1* and *Cxcl2*, and their unspliced counterparts (Fig. 4e).

Though the RNA velocity analyses of these data by both *TopicVelo* and *scVelo* suggested a quiescent-ILC3 transition Fig. 4f) and predicted the observed down-regulation of *Klf2* and *Fos* [33] during the transition (Supplementary Fig. 9a, b), only *TopicVelo* revealed the transition path of the biologically important ILC2-ILC3 trajectory or suggested a possible bi-directional quiescent-ILC2 transition (Fig. 4f). To quantitatively confirm these intertwined transitions, we computed rescaled mean first passage times (rMFPT) to different target cell populations. First, we use cells very strongly associated with the ILC3-like gene programs as target cells. The rMFPTs derived from *scVelo* show little to no variation across cells, whereas results from *TopicVelo* showed a clear distinction that suggested that, relative to transitions from other populations, the quiescent-ILC3 and ILC2-ILC3 transitions may both occur at a relatively fast timescale (Fig. 4g, Supplementary Fig. 8a, b). For quiescent-like cells as the target, both methods suggested a low likelihood of a reverse ILC3-quiescent transition; furthermore, *TopicVelo* suggested a possible ILC2-quiescent conversion (Fig. 4h, Supplementary Fig. 8c, d). For ILC2-like cells as targets, *scVelo* revealed the possible ILC2-ILC3 conversion, and both methods suggested a possible quiescent-ILC2 transition, though *TopicVelo* offered a clearer distinction between the quiescent-like cells and other origin sub-populations (Fig. 4i, Supplementary Fig. 8e, f). The discrepancies between *TopicVelo* and *scVelo* results were at least partly due to differences in velocity estimates. For example, the observed up-regulation of *Il23r*, *Il1r1*, and *Lgals3* during ILC3 response [33] was more faithfully captured by *TopicVelo* than *scVelo* (Supplementary Fig. 9c–e).

Taken together, these results suggest that the *TopicVelo* approach is effective in the analysis of immune responses, where cells may be more likely than in developmental differentiation to exhibit functional plasticity or reflect varying contributions of simultaneous, very distinct, dynamic processes.

## Discussion

RNA velocity inference has recently been improved via different machine learning techniques [18, 21–26, 62, 63]; but challenges remain. In this work, we present *TopicVelo*, a new method and framework for RNA velocity that improves on the state of the art and conceptually complements other approaches. Existing methods typically include genes based on their fit to a velocity model [8, 24–26], which makes strong assumptions about a globally determined steady state, potentially excluding genes that are informative specifically for locally dynamic processes. In contrast, by using topic modeling to discover biologically relevant gene programs or processes (“topics”) and the cells in which their activity levels are relatively high, *TopicVelo* hones in on genes that are informative for the kinetic parameters for different processes, while preventing cells that are not associated with a process from distorting its parameter estimates. To provide a global view of cell-state transitions, *TopicVelo* leverages the probabilistic topic weights to integrate process-specific transition matrices into a unified transition matrix.

*TopicVelo* infers gene-specific parameters of a transcriptional burst model by efficiently estimating the full joint distribution of unspliced and spliced gene counts given by a chemical master equation, thus explicitly accounting for higher-order moments. In contrast, the leading method *scVelo* [8] and others[18, 22, 24, 26, 62], which infer kinetic parameters based on ordinary differential equations (ODEs) from counts smoothed across cell neighborhoods in the *k*-NN graph, can distort second- or higher-order moments [15]. A recent method also incorporated a global burst model, fit via numerical gradient descent, rather than the simplex-based optimization in *TopicVelo*, though the study focused on analyzing the effects of gene-length dependent capture rates of un-spliced RNA [64]. In our analyses of real, biologically varied, single-cell datasets, we find that the transcriptional burst model enables *TopicVelo* to more accurately estimate kinetic parameters, particularly for lowly expressed genes, which can play impactful biological roles [65, 66].

A critique [16] of the *scVelo* approach notes that smoothing actually occurs at multiple stages and leads to a potentially problematic, strong dependence of the parameters, especially in the dynamical model, on the structure of the *k*-NN graph, which ideally models the underlying manifold and is visualized in the UMAP embedding. At the gene level, *TopicVelo* circumvents this issue by inferring kinetic parameters from unsmoothed counts. Furthermore, by computing a different *k*-NN for each topic, *TopicVelo* loosens the coupling between the transition matrix and UMAP embedding. While *TopicVelo*, like *scVelo*, uses the inferred velocity matrix and a matrix of differences of smoothed spliced counts to compute transition probabilities, the *TopicVelo* framework also naturally permits (noisier) transition probabilities to be computed from differences of unsmoothed counts.

Using its dissection-then-integration approach, *TopicVelo* inferred robust, accurate dynamics in complex systems, including plastic immune responses and multi-furcating differentiation, without requiring multiple time points or the support of metabolic labeling. The combination of topic modeling with a steady-state transcriptional model may allow *TopicVelo* to implicitly handle some non-steady state contexts. Future challenges include developing methods that merge the advantages of *TopicVelo* with other recent, complementary advances, such as incorporation of more sophisticated topic models [67], transcriptional models (e.g., [68]), improved transcript quantification [64, 69], a Bayesian deep generative framework for quantifying statistical uncertainty, which was developed for ODE velocity models [22, 62, 63], improvements in robustness by post-processing noisy velocity vectors using representation learning[23, 70], and multi-omic data and models [17, 19]. Another set of challenges is the interpretation of RNA velocity data. Traditional approaches heavily rely on streamline visualizations and pseudotime, which may be inadequate or misleading. In the vein of our application of fundamental Markovian techniques to quantitatively assess transition matrices, future work may borrow ideas from nonequilibrium statistical mechanics and relevant sampling frameworks, potentially leading to more reliable tools to provide mechanistic insights into cell state transitions. We believe *TopicVelo* provides a framework for developing more sophis-ticated RNA velocity methods, while serving as a valuable biological tool for inferring accurate simultaneous dynamics of interpretable gene programs and cell state transitions in diverse systems.

## Methods

### Topic modeling and differential expression analysis

We use previous work on topic model inference [37, 38, 71]. First, we use the tomotopy Python package [71] to efficiently infer topic models for a range of values of *K*, the number of topics. After evaluating those results to select a final value for *K*, we use the FastTopics R package [38] to infer the final model and compute topic-specific differentially expressed genes.

In the probabilistic topic model for scRNA-seq data, for cell *i*, *x*_*i*_ = (*x*_*i*1_, …, *x*_*iM*_) is drawn from a multinomial distribution

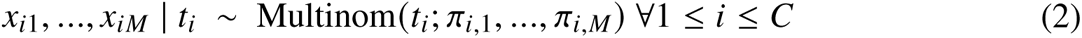

where *C* is the number of cells, *M* is the number of genes, *x*_*im*_ is number of mRNA transcripts for gene *m* in cell *i*, and 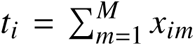 is the total number of transcripts in cell *i*. The multinomial probabilities are

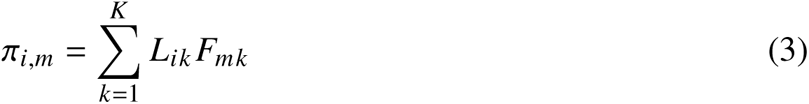

where *K* is the user-specified number of topics; 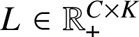 is the cell topic weight matrix, and *L*_*ik*_ is the probability of topic *k* in cell 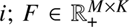 is the gene topic weight matrix, and *F*_*mk*_ is the weight of gene *m* in topic *k*.

For a given *K*, we exploit the equivalence of the maximum likelihood estimates for Poisson non-negative matrix factorization (NMF) and the multinomial topic model [38]. The negative of the log-likelihood of the Poisson NMF [72] for cell *i* and gene *m* is:

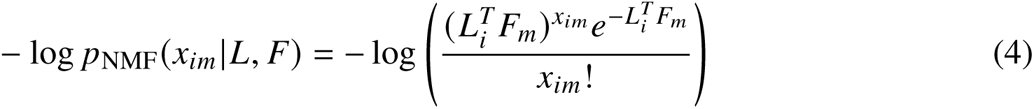

After discarding the terms that are not related to *L* and *F* and summing over all cells and genes, we arrive at a suitable loss function [38]:

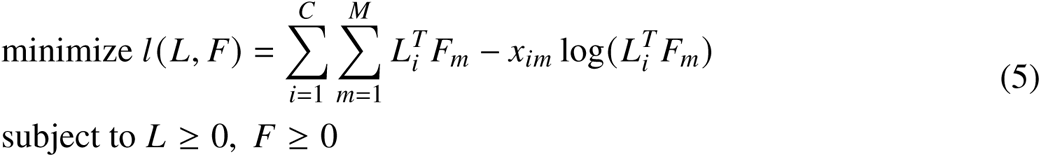

where *L*_*i*_ and *F*_*m*_ are the column vectors (of size *K*) containing row *i* of *L* and row *m* of *F*. In other words, the optimal *L* and *F* are fitted such that, accounting for the heterogeneity in cells over the topics and the contributions of individual genes to each topic, the input count matrix should be recovered on expectation.

Since each transcript count is generated by a Poisson model, the differentially expressed genes can be identified by computing the log fold change (LFC) of each gene in topic *k* [38], defined as

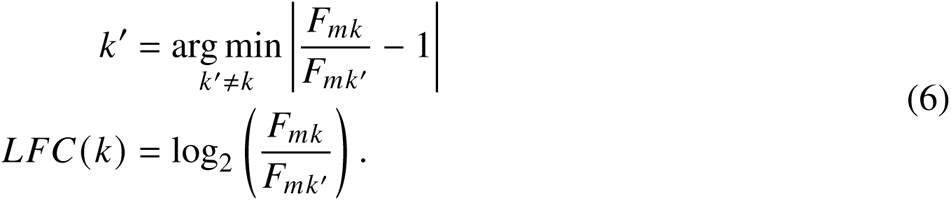

The posterior distribution of the LFC and local false sign rate (lfsr) [73] are then estimated with MCMC and stabilized with adaptive shrinkage.

For an optimized value of *K*, the above procedures were performed using FastTopics [38] as follows:

t o p i c m o d e l f i t <− f i t t o p i c m o d e l (c o u n t m a t r i x, k=K)

d e r e s u l t s <− d e a n a l y s i s (t o p i c m o d e l f i t, c o u n t m a t r i x)

where the input count matrix is constructed by stacking the raw spliced count matrix and the raw unspliced count matrix for top 2000 highly variable genes.

### Topic modeling evaluation metrics

To estimate the optimal number *K* of topics, we computed established metrics [39–41] on topic models inferred using tomotopy [71] for a range of values of *K*. For each dataset, at least one of these metrics plateaued as a function of increasing values of *K*, and we selected the smallest value of *K* in the intersection of those regimes across metrics.

The first metric we considered uses average distance among topics to measure stability [39]:

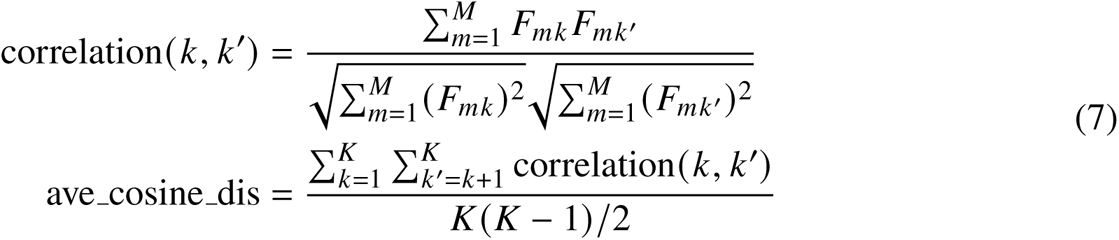

where correlation(*k*, *k*^′^) is the standard cosine distance between topics *k* and *k*^′^. A smaller ave cosine dis indicates more stability.

The second metric we considered is the information divergence between all pairs of topics [40]:

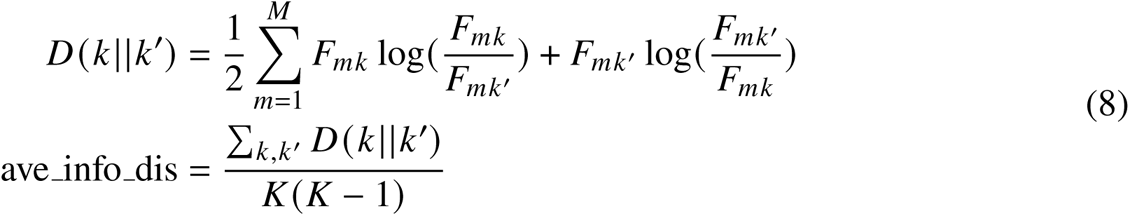

where *D*(*k* ॥ *k*^′^) is the Jensen-Shannon distance between two topics. A bigger ave info dis indicates more independence and more information in the topic model.

We also tested a few coherence measures, which are based on the point-wise mutual information (PMI) of the top-weighted or highest ranked (by log-fold change) topic-specific genes [41]:

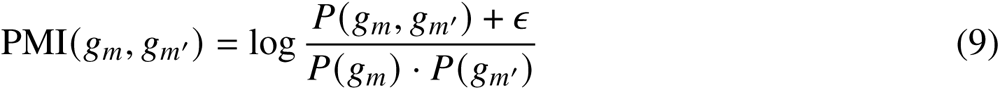

where *P*(*g*_*m*_, *g*_*m*_′) is the joint probability of observing genes *g*_*m*_ and *g*_*m*_′ in a cell, and *P*(*g*_*m*_) and *P*(*g*_*m*_′) are the marginal probabilities of observing gene *g*_*m*_ and *g*_*m*_′ in a cell, respectively; *∊* is a small number (e.g. 10^−12^) to prevent the PMI from reaching 0. For the top *N* genes (either topic-specific or highest weighted), the *UCI coherence* is calculated as:

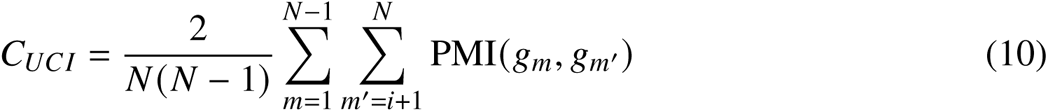

Small values of |*C*_*UCI*_ | indicate higher topic coherence and higher probability that the top genes are co-expressed.

Finally, to prevent overfitting, we also consider the Akaike information criterion (AIC) and the Bayesian information criterion (BIC):

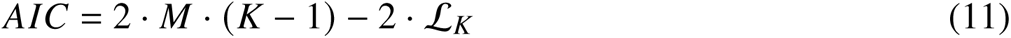

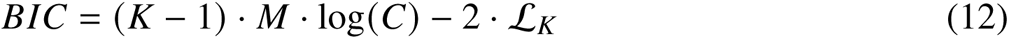

where *ℒ*_*K*_ is the log-likelihood of the model for *K* topics.

In addition, interpretability, i.e. a reasonable number of potentially biologically meaningful differentially expressed genes, is another important criterion. For most datasets, “topic-specific genes“ were selected from the differentially expressed genes for downstream analysis (e.g., RNA velocity) if, for either the spliced or unspliced form, the lfsr is at most 0.001 and the LFC is at least 0.5 in absolute value. This criterion is a very conservative estimate of differential expression and, in practice, produces 50–250 topic genes for each topic.

### RNA velocity parameter estimation via the one-state model

The one-state transcription model is governed by the following master equation:

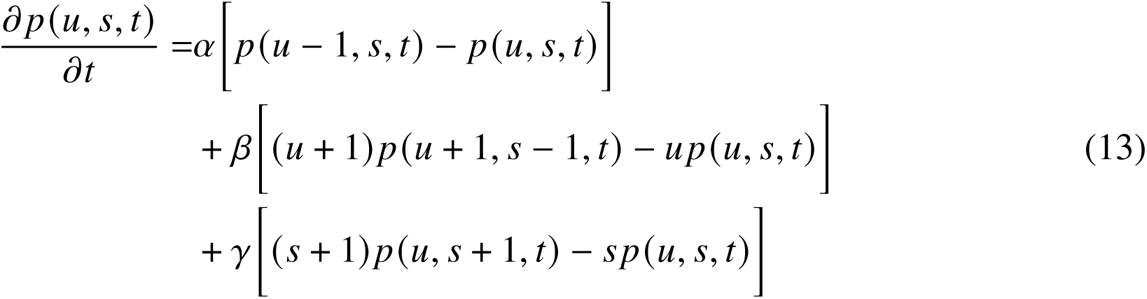

in which *α* is the rate of transcription, *β* is the splicing rate, and *γ* is the degradation rate. The steady-state distribution when *β* ≠ *γ* is the product of two independent Poisson distributions for *u* and *s* respectively [74]:

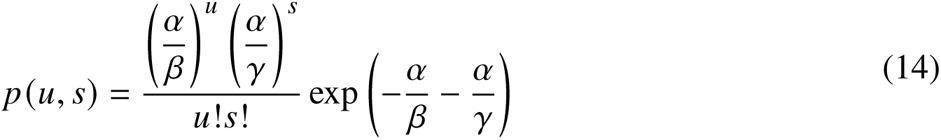

Then the log likelihood for observing *C* cells at steady state with unspliced and spliced counts 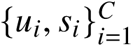 conditioned on a set of kinetic parameters is:

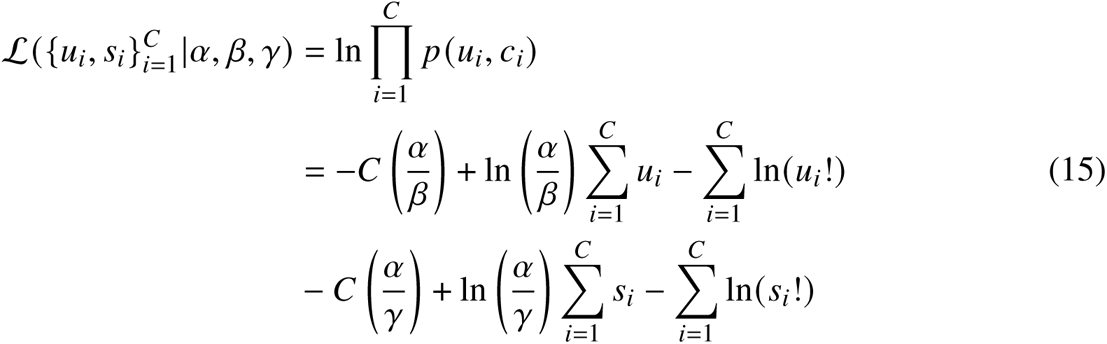

The maximum likelihood estimate of *γ*/*β* is:

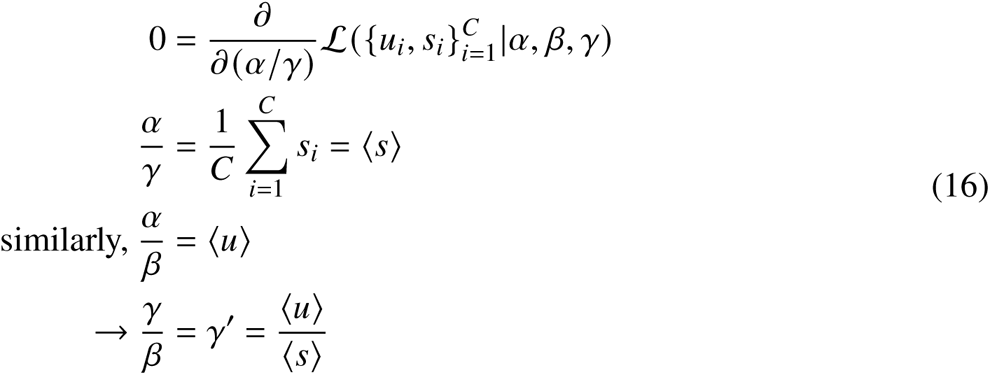

where ⟨·⟩ denotes expectation, and ⟨*s*⟩ and ⟨*u*⟩ are the average abundance of *u* and *s* over all cells in steady-state.

We note that, rather than using this analytical estimate on unsmoothed counts, *scVelo* in *stochastic* mode actually computes the moments for each cell using a *k*-NN graph. For each gene, a generalized least squares is performed by solving a system of linear equations involving the first and second moments for the cells in steady state (top right corner of the (*u*, *s*) phase plot) [8]. Though this does not agree with the analytical estimate exactly, the deviation is small in practice for unsmoothed data, since the first and second moments are time-invariant under the steady-state assumption. However, different choices for constructing the *k*-NN graph can affect parameter estimates in unexpected ways [15, 16].

### RNA velocity parameter estimation via the geometric burst model

To estimate the steady-state joint distributions, we implemented a Gillespie algorithm [42] to simulate the master equation (Equation 1) in Python, accelerated via Numba [75]. The burn-in period represents the time before the system converges to a steady state. For a trajectory with burn-in period *t*_burn-in_ and total simulation time *t*_total_, the probability *p* (*u*, *s*) of observing a cell with *u* unspliced mRNA and *s* spliced mRNA for a given gene in the steady state is

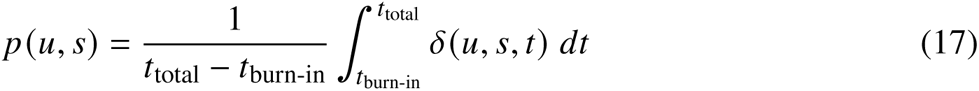

where *δ*(*u*, *s*, *t*) = 1 if the cell has *u* unspliced counts and *s* spliced counts at time *t*, and *δ*(*u*, *s*, *t*) = 0 otherwise.

To infer the kinetic parameters governing the dynamics, we initialize the parameters with the method of moments, which was previously derived [31, 64]:

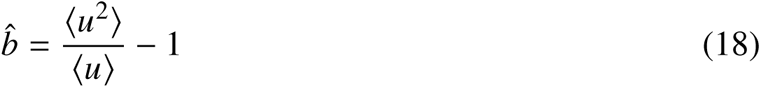

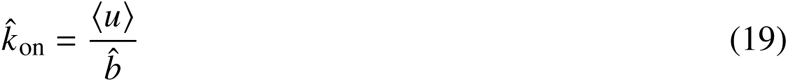

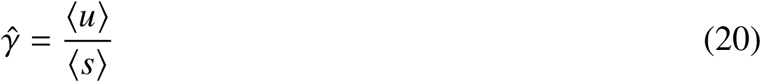

where the moments are estimated from the observed distribution. Then to find the optimal kinetic parameters, the KL divergence is minimized using the Nelder-Mead algorithm implemented in SciPy [43]. In some cases, the method of moments estimate is a local minimum that is close to the global minimum, and the optimizer can get stuck. In this case, we used 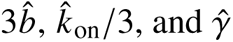 to restart the search for the global minimum. The convergence criterion was chosen to be a relative change in KL divergence between two subsequent iterations smaller than 1/1000 or reaching a maximum number of iterations.

To verify the correctness of this estimation approach, we compared the simulated joint distribution for parameters *k*_on_ = 0.5, *b* = 5, *γ* = 3 with the joint distribution simulated from the inferred parameters; the two distributions are nearly identical (Supplementary Fig. 1a). To visualize the path of the optimization, we plotted it on the KL divergence landscape of *k*_on_ versus *b* for *γ* fixed at 0.3, and the KL divergence landscape of *γ* versus *b* with for *k*_on_ fixed at 0.5; we observed that the optimizer ended very close to the ground truth (Supplementary Fig. 1b, c).

We aimed to find choices for the number of simulation steps (or reactions) and the maximum number of iterations for the Nelder-Mead optimization that perform well in this inference scheme. We considered a total of 27 parameter combinations across different dynamical regimes in which average mRNA abundances vary over the span of several orders of magnitudes (Supplementary Fig. 1d). First, we fixed the number of simulation steps at 5 × 10^5^ and analyzed how different choices for the maximum number of optimization iterations affect the performance. For each choice, we simulated 10 replicates of the 27 combinations and computed the average KL divergence within each replicate (Supplementary Fig. 1e). We observed that the performance stopped improving when more than 50 iterations were used. Similarly, we fixed the maximum number of iterations at 50 and examined how different choices for the number of simulation steps affected inference. We observed that the performance improvement became negligible when more than 5 × 10^5^ steps were used (Supplementary Fig. 1f). Therefore, we chose 5 × 10^5^ as the number of reactions and 50 as the maximum number iterations for parameter inference. In both settings, the bimodality of the average KL divergence that we observed may be due to the optimizers getting stuck in local minima. To ameliorate this effect in the analysis of real datasets, we perform 5 independent optimizations for each gene and, for downstream analysis, use the set of parameters that corresponds to the lowest KL-divergence.

In the scRNA-seq data applications, the joint distribution of spliced and unspliced counts was typically computed from the size-normalized data (not on the log scale), rounded to the nearest integer. Under the steady-state assumption, a time-invariant splicing rate *β* = 1 was assumed for all genes; *k*_on_, *b*, and *γ* were estimated for each gene. To illustrate the parameter inference scheme on real data, we used the observed distributions of *Grin2b* from the granule mature cells in the dentate gyrus dataset [76], which serve as a proxy for steady state since these cells are terminally differentiated. By capturing the diffusiveness and low expression regimes of the distribution more accurately, the geometric burst model recovers a joint distribution that is closer than the one inferred via the one-state model to the observed distribution (Supplementary Fig. 1g). The inferred parameters from the burst model for *Grin2b* are located within a regime of low KL divergence as shown on the KL divergence landscapes (Supplementary Fig. 1h). We performed the analogous analysis for the gene *Btbd9* and observed similar results (Supplementary Fig. 1i, j).

### Determination of topic-associated cells

For each topic within a given dataset, topic-associated cells are defined as cells above a certain topic weight. Kinetic parameters for topic-specific genes are inferred from topic-associated cells, which in this model are assumed to represent a topic-specific steady state. In the *scVelo* implementation, the up-regulation and down-regulation steady states for a given gene are modeled as the top right corner and the bottom left corner of the phase plot, respectively. The exact determination is dependent on arbitrary expression thresholds; the default setting uses the 5th and 95th percentiles. Instead of assuming each gene has its own set of steady-state cells, *TopicVelo* uses topic association as a criterion for choosing topic-specific steady state cells, which tends to be more robust and biologically meaningful because the genes in the topic-specific gene programs have correlated expression patterns.

While one approach for choosing a topic weight threshold is to associate each cell with the topic in which it has the highest weight, which discretely clusters the cells, this has a number of drawbacks: (1) a cell may have relevant information about a topic in which it participates but for which it does not have the highest weight; (2) by the same token, the cells assigned this way to a topic may not capture the full dynamic range of an associated process; and (3) for the purpose of computing transitions, this approach is problematic because there is no potential for transitions between cells assigned to different topics.

In general, we used the following procedure to identify a reasonable range for the choice of topic weight threshold. For a given topic *k*, we denote the set of cells with topic-*k* weights above the *n*^*th*^-percentile as 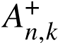, and the set of cells with topic-*k* weights below or equal to the *n*^*t*ℎ^-percentile 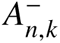. Note that 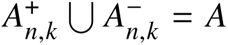 where *A* is the set of all cells. For integers *n* from 1 to 99, we compute what we call an *average rescaled KL divergence*, denoted by 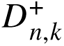 as follows: for each topic-specific gene, we compute the KL divergence of the joint *u*-*s* distribution of 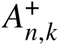 to that of *A*, and rescale the divergence to [0, 1]; then we average the rescaled KL divergences over the genes. We perform an analogous procedure to compute 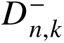, the average rescaled KL divergence for the distribution from 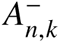 to that of 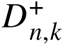 *n* approaches 0. We observed a sharp approachs 0 as decline in 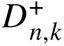 at a relatively large value of *n*, which we denote by 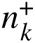. If the topic weight threshold is chosen in the regime 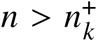, the full dynamic range of topic-associated process is not properly accounted for. Similarly, 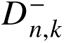 approaches 0 as *n* approaches 100, and a sharp decline in 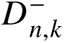 is observed for a relatively small value of *n* denoted by 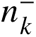. Topic weight thresholds in the regime 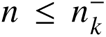 risk including cells not meaningfully associated with the topic-associated process. The interval 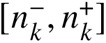 is a natural and simple heuristic for the range of suitable thresholds for topic *k*. For the majority of topics and datasets, we observed 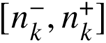 to be a range in which both 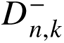 and 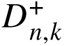 were relatively flat, though in other cases, this range was observed be around [75, 95], and these topics often corresponded to a rare cell type or when the process is very distinct.

### Construction of topic-specific transition matrices

While we use unsmoothed counts for kinetic parameter inference, we compute the transition flows on smoothed counts to remove noise in the visualization. However, we did not observe significant distortions in the overall trends using smoothed versus unsmoothed counts. For cell *i* and gene *m*, the first moments 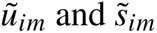 represent the smoothed counts, computed as the number of unspliced and spliced transcripts, respectively, averaged over the cells in the neighborhood of *i* in the NN (for 30 nearest neighbors) graph, computed from the top 30 PCs of the global principal components (PC) analysis of the log-normalized spliced expression matrix.

The velocity vector for cell *i* associated to topic *k* is 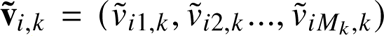, for topic-specific velocity vector 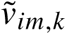 defined as 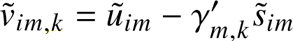 for gene *m*, where *M*_*k*_ is the number of topic-specific genes, and 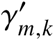. is the topic-specific degradation rate for gene *m.* Across small neighborhoods in the NN graph, the first moments of the smoothed data are not as distorted as higher-order moments, and the velocity 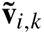 is a reasonable smoothed approximation.

Then a cosine similarity between the velocity vectors and the differences in spliced expression can be computed, as previously done [8]:

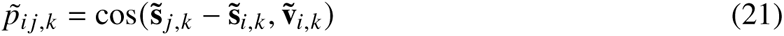

where 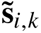 is the vector of smoothed spliced counts in cell *i* for topic-*k* specific genes.

For each topic *k*, a topic-specific NN graph is constructed on just the topic-associated cells using the top 30 PCs of the global PCA as distances. The topic-specific transition probability *p*_*i* *j*,*k*_ from cell *i* to cell *j* for topic *k* is obtained by applying an exponential kernel to the cosine similarities over the set *N*_*k*_ (*i*) of cells in the topic-specific neighborhood of cell *i*:

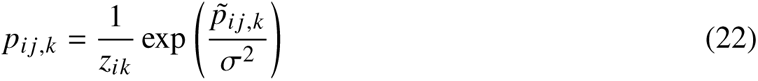

where *σ* is the kernel width parameter and 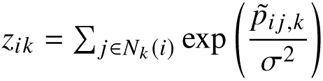 is the normalization factor.

### Integration of process-specific dynamics

Because the topic-associated cells and global set of cells may have different indices, we switch to using *c* to denote the identity of a cell instead of using its index. To compute the global transition matrix, we first renormalize the topic weights 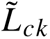 over just the topics that cell *c* is associated to:

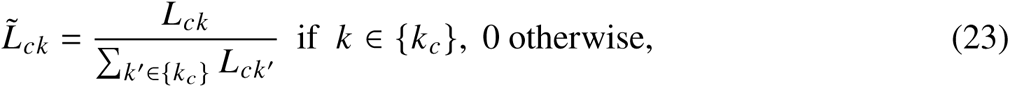

where {*k*_*c*_} is the set of topics associated to cell *c*.

The global probability of a transition from cell *c* to *c*^′^ is computed as

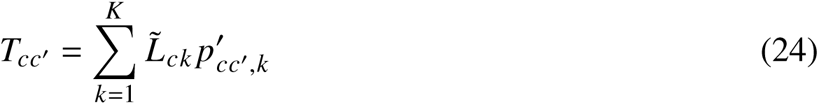

where 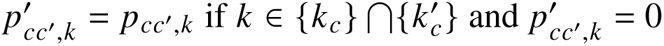 otherwise.

### RNA velocity evaluation metrics

We use the stationary distribution to assess the overall directionality of the transition matrix. We use the mean first passage time (MFPT) to evaluate the short-term dynamics and the directionality of transient conversions. Without loss of generality, suppose the integrated transition matrix *T* is irreducible. By construction, *T* is positive-recurrent and aperiodic. The stationary distribution *π* is the solution to the eigenvalue problem *π*^*T*^ = *π*^*T*^*T* . The MFPT matrix *M* (where element *M*_*i*, *j*_ is the MFPT from state *i* to *j*) is the solution to the following matrix equation:

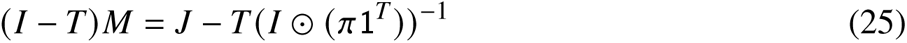

where *I* is the identity matrix and *J* is a matrix of all ones. For a set of target cells *C*_*a*_, the vector of MFPTs to *C*_*a*_, denoted as *M*_*Ca*_ is

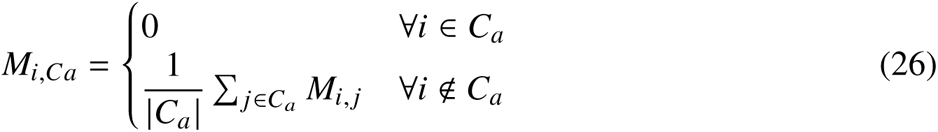

To highlight trends beyond absolute magnitude, for each velocity method, for each *C*_*a*_, we rescale *M*_*Ca*_ by the median of nonzero elements in *M*_*Ca*_ to obtain the rescaled mean first passage time (rMFPT). If extremely distinct populations are contained within a given dataset, *T* may be reducible (i.e., *T* contains multiple disconnected components), in which case the stationary distribution and MFPT must be analyzed within individual irreducible components. This was not an issue in the datasets analyzed here.

### Preprocessing of scRNA-seq datasets

For each dataset, genes were filtered so that there are at least 20 cells that have both spliced and unspliced mRNA transcripts for each gene. Counts were size-normalized to the median total counts, including spliced and unspliced transcripts.

A principal components analysis was performed on the log-normalized spliced counts matrix using the top 2000 highly variable genes. From the top 30 principal components, a *k*-nearestneighbor (*k*-NN) graph was constructed (using the default of *k* = 30). (We use the standard parameter terminology, but the *k* in the definition of the *k*-NN is completely independent of the parameter *k* in the topic model.)

Then the first and second moments of each cell were estimated over the *k*-NN graph. The above procedures were performed via *scVelo* [8]:

sc Velo . pp . f i l t e r a n d n o r m a l i z e (a d a t a, m i n s h a r e d c o u n t s = 20) sc Velo . pp . moments (a d a t a, n p c s = 30, n n e i g h b o r s = 30)

### Analysis of the human hematopoiesis scNT-seq data

This dataset contains count matrices with or without metabolic labels. We focused our analysis on the latter. We used default settings for the *scVelo* stochastic and dynamical models to infer velocities and obtained streamline embeddings. The dynamical model gave streamline embeddings more consistent with biological expectation. We then applied topic modeling with 8 topics and identified topic-specific genes. We removed topic-specific genes from topics 5 and 6 for downstream analysis because they are strongly associated with minichromosomal and ribosomal genes and are ubiquitously expressed. We selected topic-associated cells as those with weights above the 65^th^-percentile for each topic to infer the kinetic parameters and global transition matrix. We compared topic-specific velocities to the global velocity inferred by the *scVelo* dynamical model.

### Analysis of the mouse gastrulation data

After standard preprocessing, we applied the *scVelo* stochastic model, rather than the dynamical model, which is more prone to wrongly inferring transcriptional boosting as down-regulation [12]. We performed topic modeling with 2 topics, which resulted in a large number (1,961) of topic-specific genes. To focus the analysis on the genes with the best signal given the low unspliced/spliced ratio in this dataset, we removed genes for which the ratio of the maximum of spliced counts to the maximum of unspliced counts is greater than 10 or less than 0.01, as the ratios of spliced over unspliced counts in this dataset tend to be very high. Furthermore, we observed that this dataset contains many genes that are highly expressed in specific cell subsets, which standard size-normalization steps do not handle well, potentially leading to improper assessment of the dynamics of lowly expressed genes. To avoid this possibility, we used the raw counts (rather than the size-normalized counts) to infer kinetic parameters. We used cells with topic weights above the 40^th^ percentile for each topic to infer the kinetic parameters and global transition matrix. We compared topic-specific velocities to the global velocity inferred by the *scVelo* dynamical model.

### Analysis of the human bone marrow data

After standard preprocessing, we applied the *scVelo* stochastic model. Like the mouse gastrulation data, this data contain genes with transcriptional boosting patterns that are not handled well by the dynamical model [13]. We performed topic modeling with 10 topics. We used cells with topic weights above the 65^th^-percentile for each topic to infer the kinetic parameters and to construct the global transition matrix.

### Analysis of the mouse ILCs data

This dataset contains data from five different days. We only used the data collected on day 3. To maintain comparisons with the previous analysis, we focused on the highly variable genes as determined previously by a variance stabilizing transformation [33]. We still performed gene filtering, and then performed topic modeling with 10 topics. For the global analysis, we used the 82^nd^-percentile for each topic to infer the kinetic parameters and global transition matrix. For analyzing velocities, we focused on comparing the *TopicVelo* topic-specific velocities and the global velocity from the *scVelo* dynamical model; in this dataset, the stochastic and dynamical *scVelo* models give very similar results. For the mean-first passage time analysis, the target cells were selected as the cells above 95^th^-percentile from topics 4, 6, and 9 respectively.

### Data availability

The gastrulation [14] and bone marrow [32] data are available in the *scVelo* package [8]. The human hematopoiesis scNT-seq [18] and ILCs data [33] and are available in the NCBI Gene Expression Omnibus (GEO) under accession numbers GSE193517 and GSE149622, respectively.

### Code availability

The source code, Jupyter notebooks, and R markdown files for reproducing figures and results in this paper are available for reviewers during the editing process. *TopicVelo* will be available as an open-source Python package for public use.

## Acknowledgements

We thank Hope Anderson for helpful discussions about interpreting the topic modeling results of the immune datasets. We thank Hanna Hieromnimon for help in the design and creation of the overview figure. S.V. and C.F.G. were supported by the NIH NIGMS Award R35GM147400.

## Author contributions

C.F.G, S.V., and S.J.R. conceived the overall project. S.V. and S.J.R. supervised the research. C.F.G., S.V., and S.J.R. conceived using the burst model. C.F.G and S.J.R conceived the topic modeling approach. C.F.G. developed the algorithm, and implemented and tested *TopicVelo*. C.F.G. and S.J.R. analyzed the data. C.F.G, S.V, and S.J.R discussed and interpreted the results, and wrote the manuscript. All authors read and approved the final manuscript.

## Ethics declarations

The authors declare no competing interests.

---

**Supplementary Figure 1:**
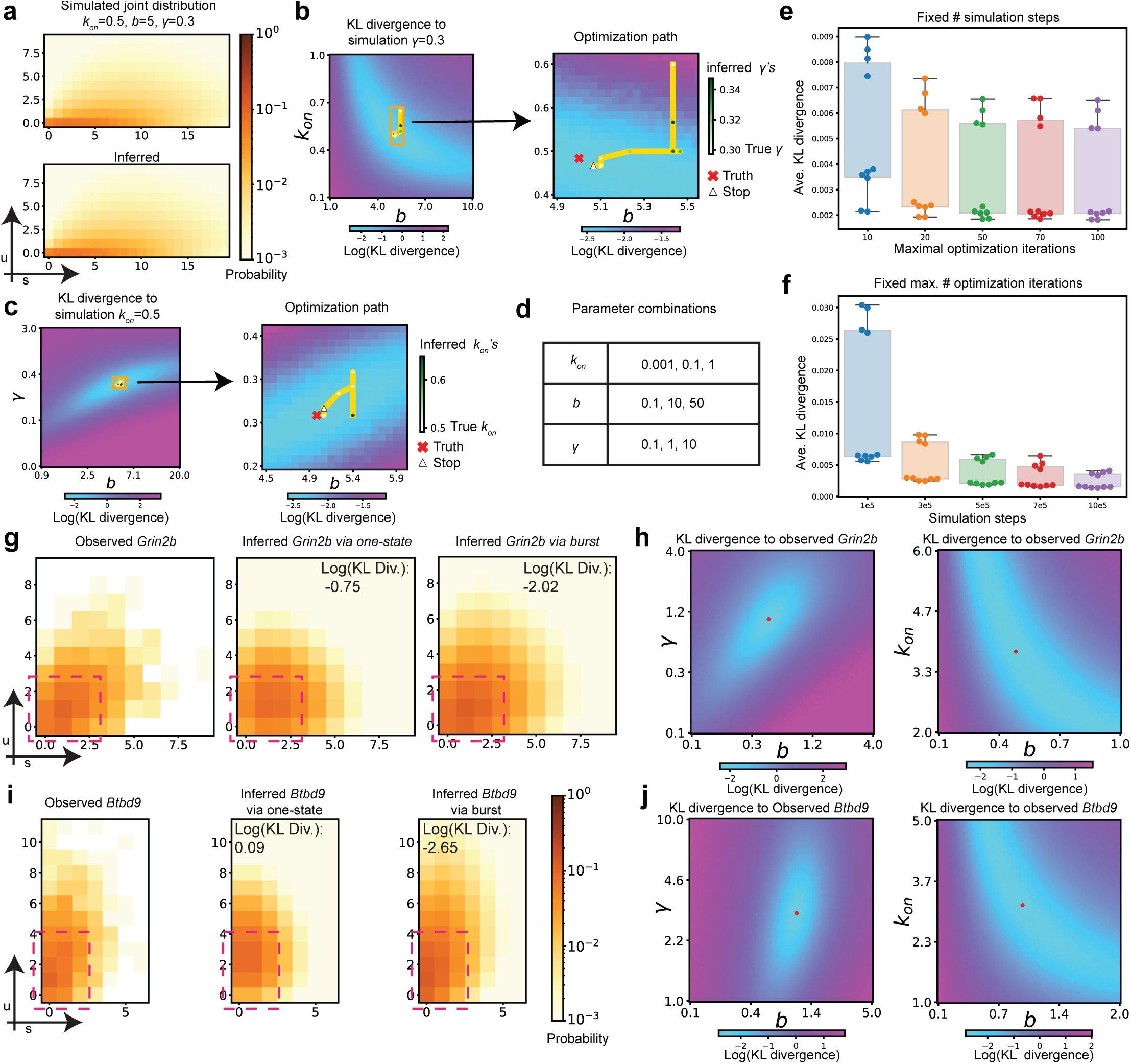
The geometric burst model more accurately recovers experimental distributions than the one-state model. **a**, An example joint distribution of spliced (*s*) and unspliced (*u*) transcript counts, as simulated by the Gillespie algorithm for the geometric burst model with fixed parameters *k*_on_ = 0.5, *b* = 5, and *γ* = 0.3 (top), and with maximum-likelihood estimates (MLEs) inferred from the simulated data (bottom). **b,** Log KL divergence (color) landscape for *γ* = 0.3 over a range of values of *b* and *k*_on_, with close-up (right) of restricted range (orange box, left). True parameter values marked by red cross; optimization path (yellow) shown across iterations (points, colored by inferred *γ* value) to end point (triangle). **c,** Analogous to b, for *k*_on_ = 0.5 and varying *b* and *γ*. **d,** Table of 27 parameter combinations used to assess effects of the number of simulation steps and maximum number of optimization iterations. **e,** For a fixed number (5 · 10^5^) of simulation steps and varying maximum number of iterations (x axis, color), bar plots show the average KL divergence across the 27 parameter combinations in d, for 10 replicates (points). **f,** Analogous to e, for a fixed maximum number (50) of optimization iterations. **g,** The joint distribution of the gene *Grin2b* in the granule mature cells in the dentate gyrus dataset [76], as observed (left), computed from MLEs for the one-state model (middle), and simulated from the MLEs for the geometric burst model (right), annotated by log KL divergence from the observed. The burst model better matches the observed values in the region where probability mass is concentrated (pink dashed box). **h,** For the burst model, plots show the log KL divergence to the observed distributions of *Grin2b* for *b* versus *γ* with *k*_on_ fixed to the MLE (left), and *b* versus *k*_on_ with *γ* fixed to the MLE (right), with optimizer end points indicated (red dots). **i, j,** Analogous to g, h, but for *Btbd9*. The log of KL divergence is base 10.

**Supplementary Figure 2:**
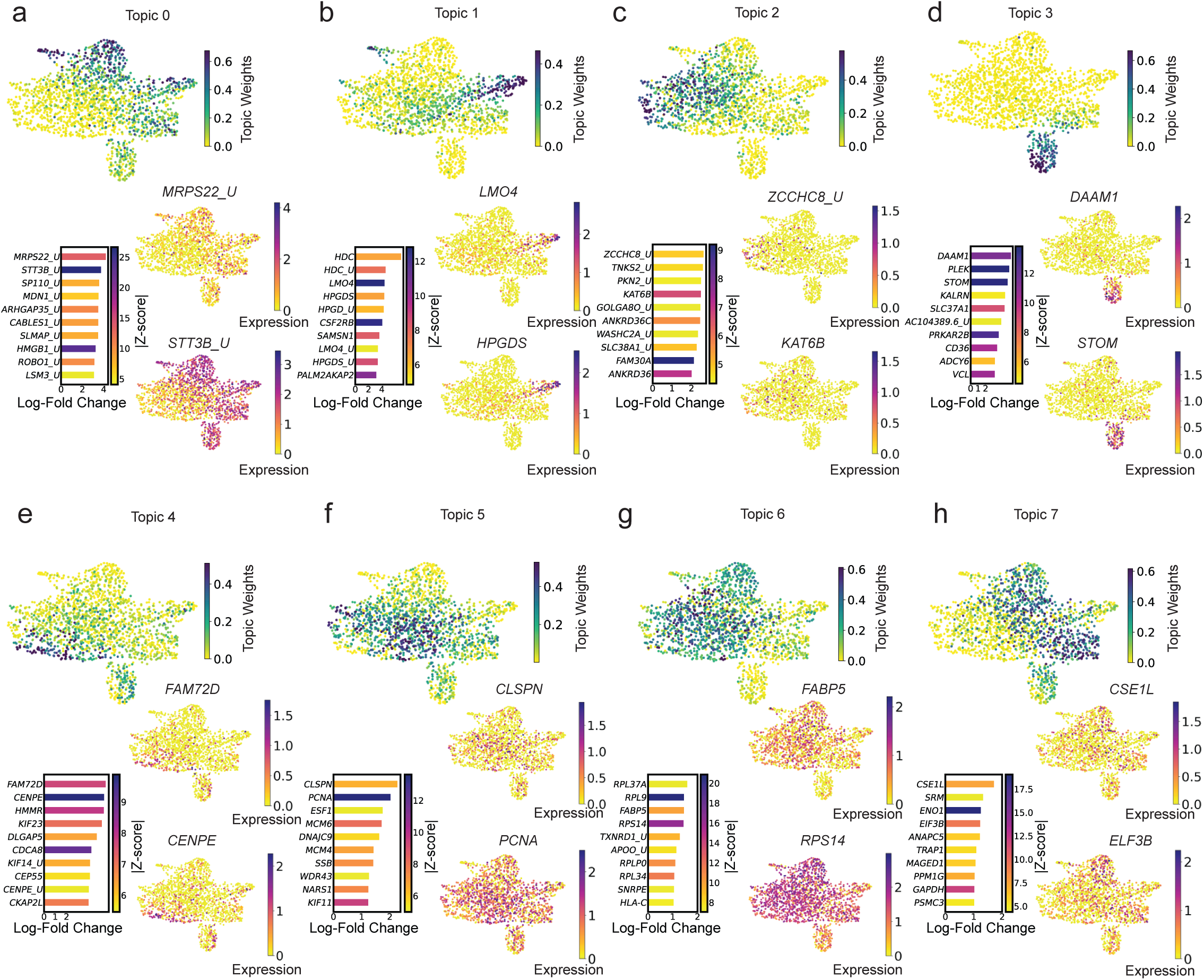
Topic modeling analysis of scNT-seq data from human hematopoiesis. **a**, For topic 0, UMAP plots shows cells colored by topic weights (top) and by log-normalized expression of topic-specific genes (bottom right); bar plot (bottom left) shows top 10 topic-specific genes ranked by log-fold change (x axis) and colored by absolute value of z-score; *’ U’* indicates unspliced transcripts. **b–h,** Analogous to a, for topics 1–7, respectively.

**Supplementary Figure 3:**
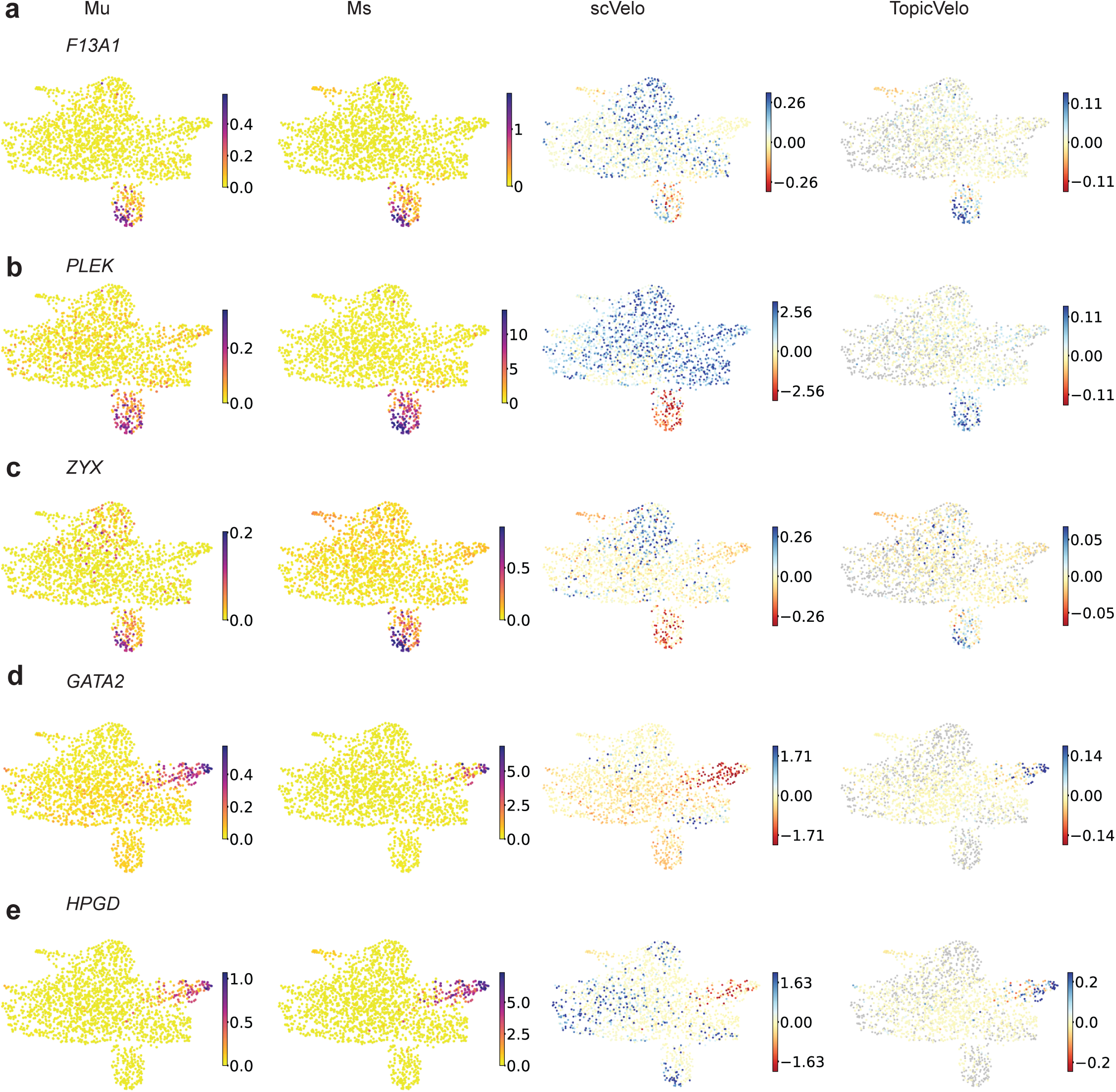
*TopicVelo* recovers more biologically plausible velocity estimates than those of *scVelo* for the scNT-seq data. **a–c**, Analysis of topic-3 specific genes. UMAP plots colored by smoothed size-normalized counts of unspliced (Mu) (far left) and spliced (Ms) (middle left) transcripts, and by velocities inferred by *scVelo* (middle right) and *TopicVelo* (far right), for the genes *F13A1* (a), *PLEK* (b), and *ZYX* (c). d, e, Analysis of topic-1 specific genes *GATA2* and *HPGD*, analogous to a–c.

**Supplementary Figure 4:**
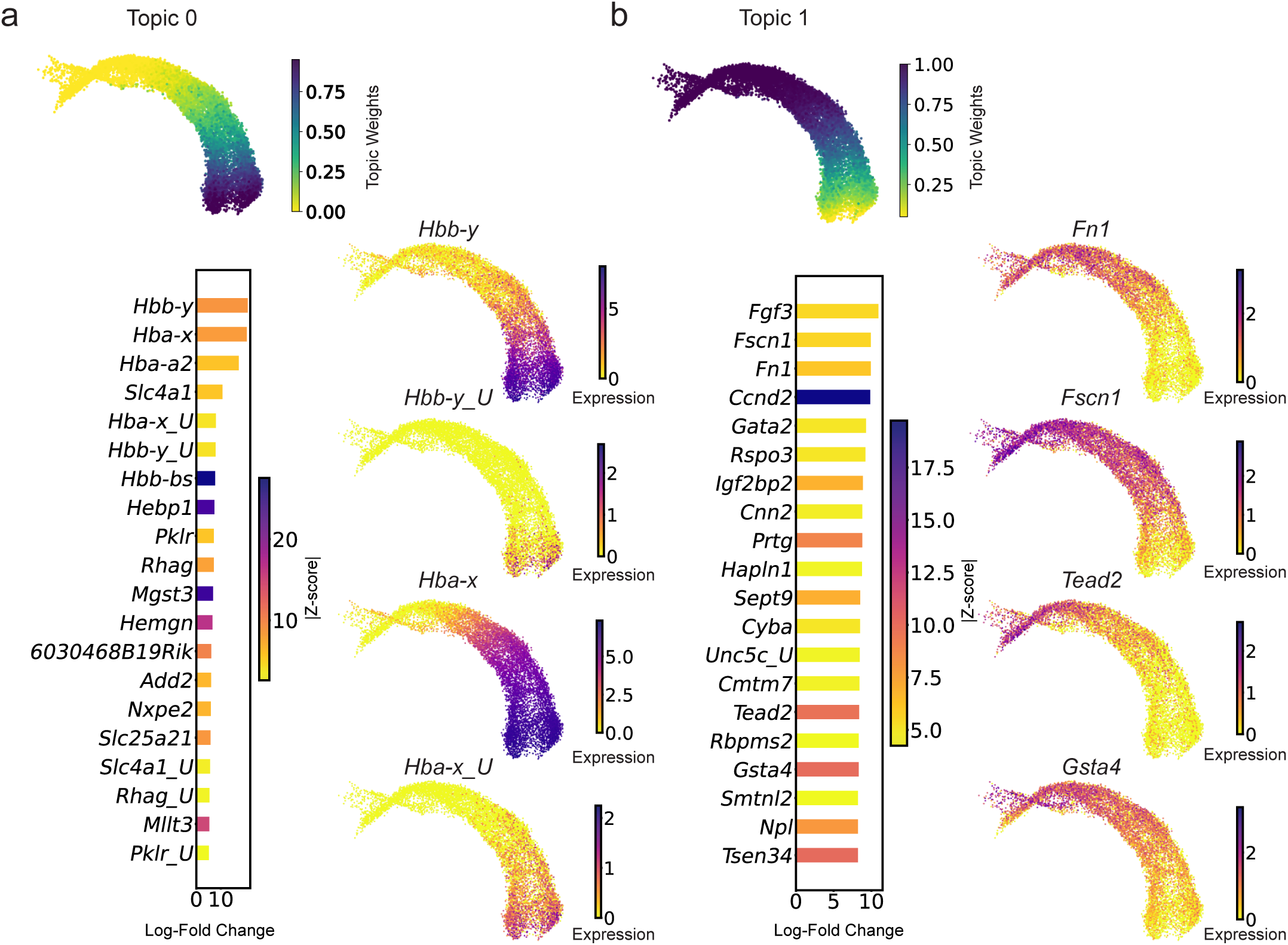
Topic modeling of the gastrulation data revels key genes underlying the differentiation of blood progenitors to erythroid. **a**, For topic 0, UMAP shows cells colored by topic weights (top) and by log-normalized expression of topic-specific genes (bottom right); bar plot (bottom left) shows top 20 topic-specific genes ranked by log-fold change (x axis) and colored by absolute value of z-score; *’ U’* indicates unspliced transcripts. **b,** Analogous to a, for topic 1.

**Supplementary Figure 5:**
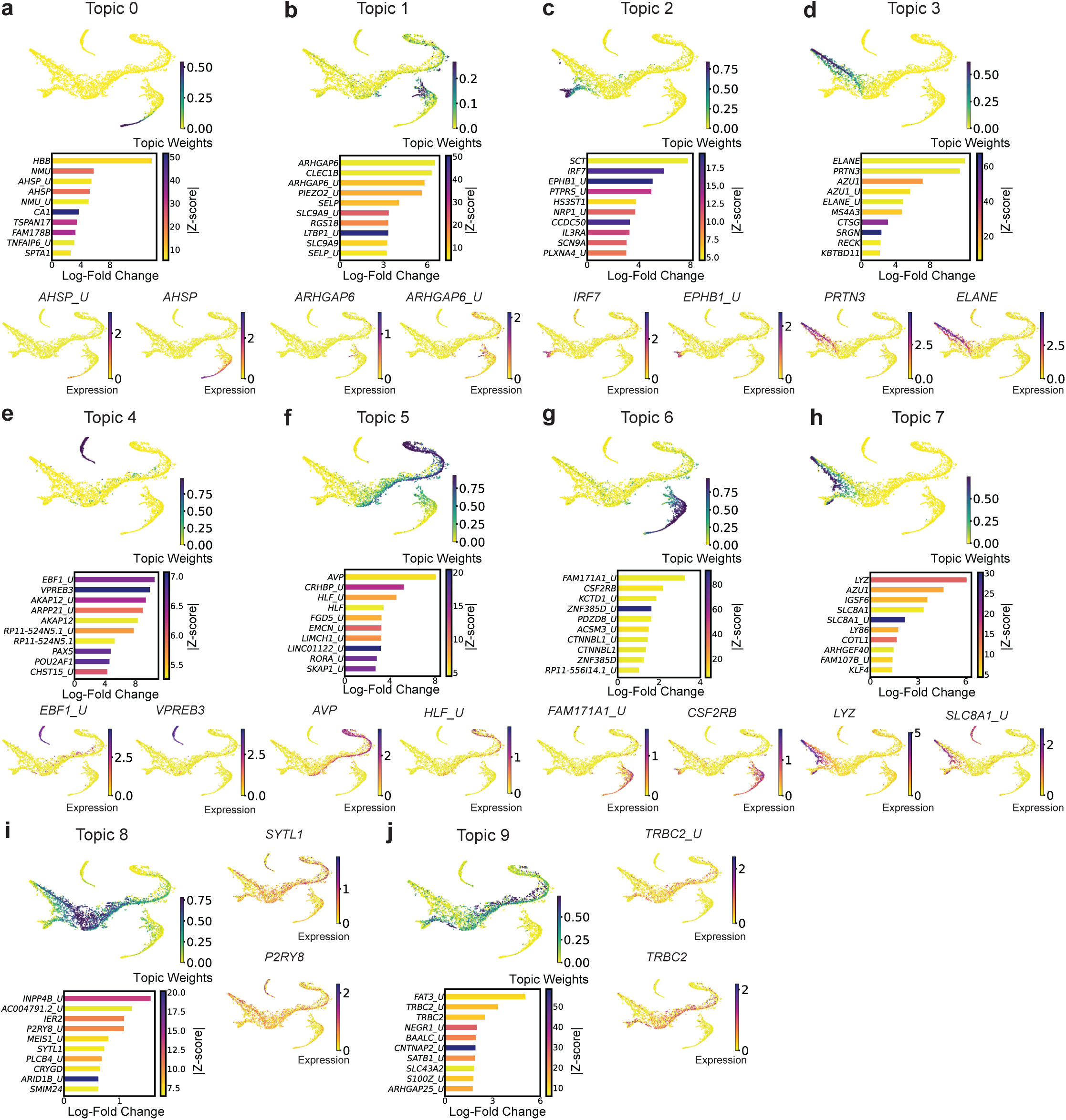
Topic modeling of the human bone marrow data provides insights into the different stages of differentiation along all lineages. **a**, For topic 0, *t*-SNE plots shows cells colored by topic weights (top) and by log-normalized expression of topic-specific genes (bottom); bar plot (middle) shows top 10 topic-specific genes ranked by log-fold change (x axis) and colored by absolute value of z-score; *’ U’* indicates unspliced transcripts. **b–j,** Analogous to a, for topics 1–9, respectively. Topics are generally associated with annotated stages of development: topic 0 (mature erythroid), 1 (megakaryocyte), 2 (dendritic cells), 3 (one lineage of monocytes), 4 (common lymphoid progenitors), 5 (hematopoietic stem cells), 6 (erythroid), 7 (another lineage of monocytes), 8 (precursors to monocytes), 9 (hematopoietic stem cells and precursors to dendritic cells).

**Supplementary Figure 6:**
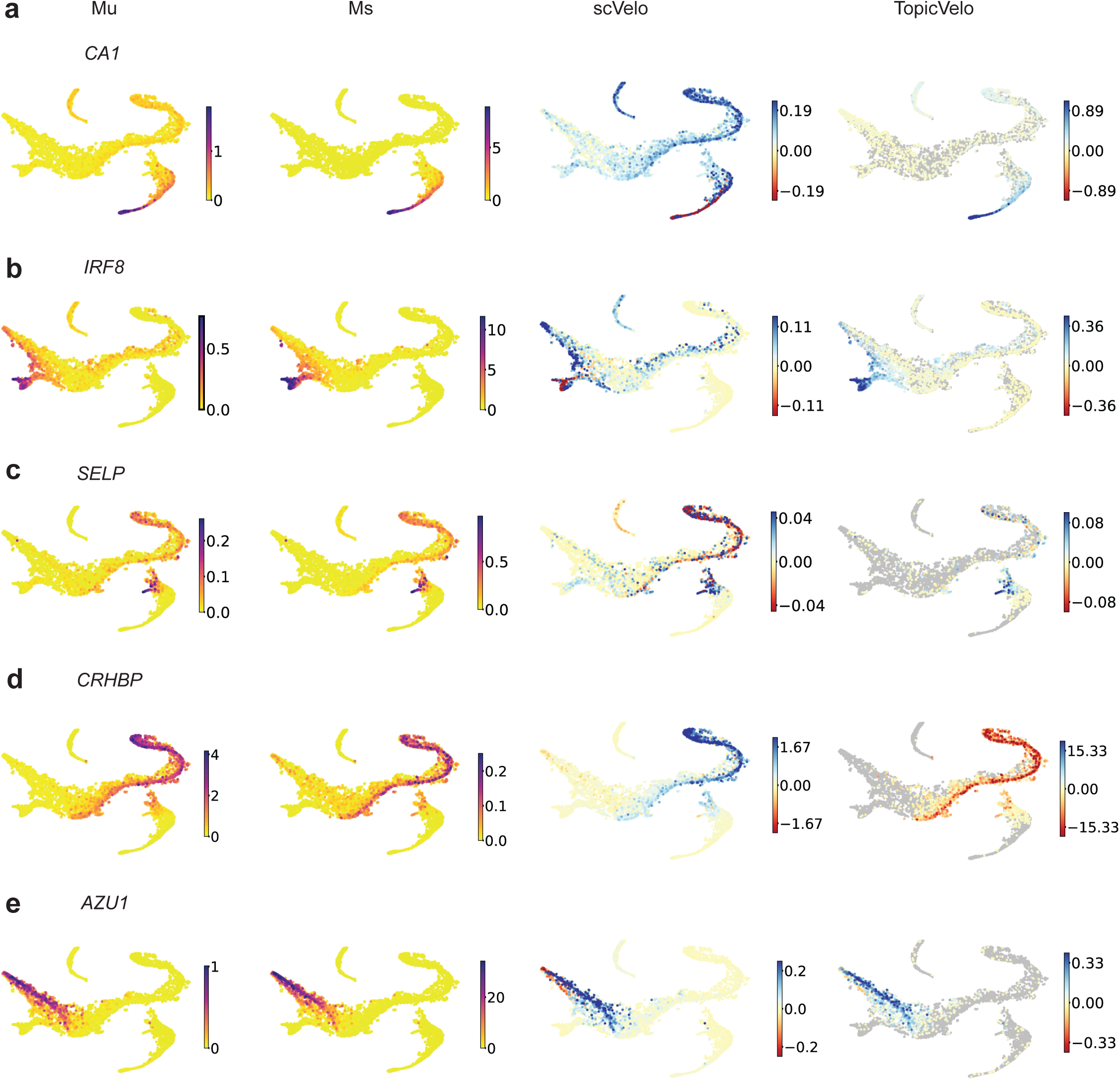
*TopicVelo* recovers better biologically supported velocities than *scVelo* for the human bone marrow data. **a**, For topic-0 specific gene *CA1*, UMAP plots are colored by smoothed size-normalized counts of unspliced (Mu) (far left) and spliced (Ms) (middle left) transcripts, and by velocities inferred by *scVelo* (middle right) and *TopicVelo* (far right). **b–e,** Analogous to a, for topic-2 specific gene *IRF8* (Supplementary Table 1), topic-1 specific gene *SELP*, topic-5 specific gene *CRHBP*, and topic-7 specific gene *AZU1*, respectively.

**Supplementary Figure 7:**
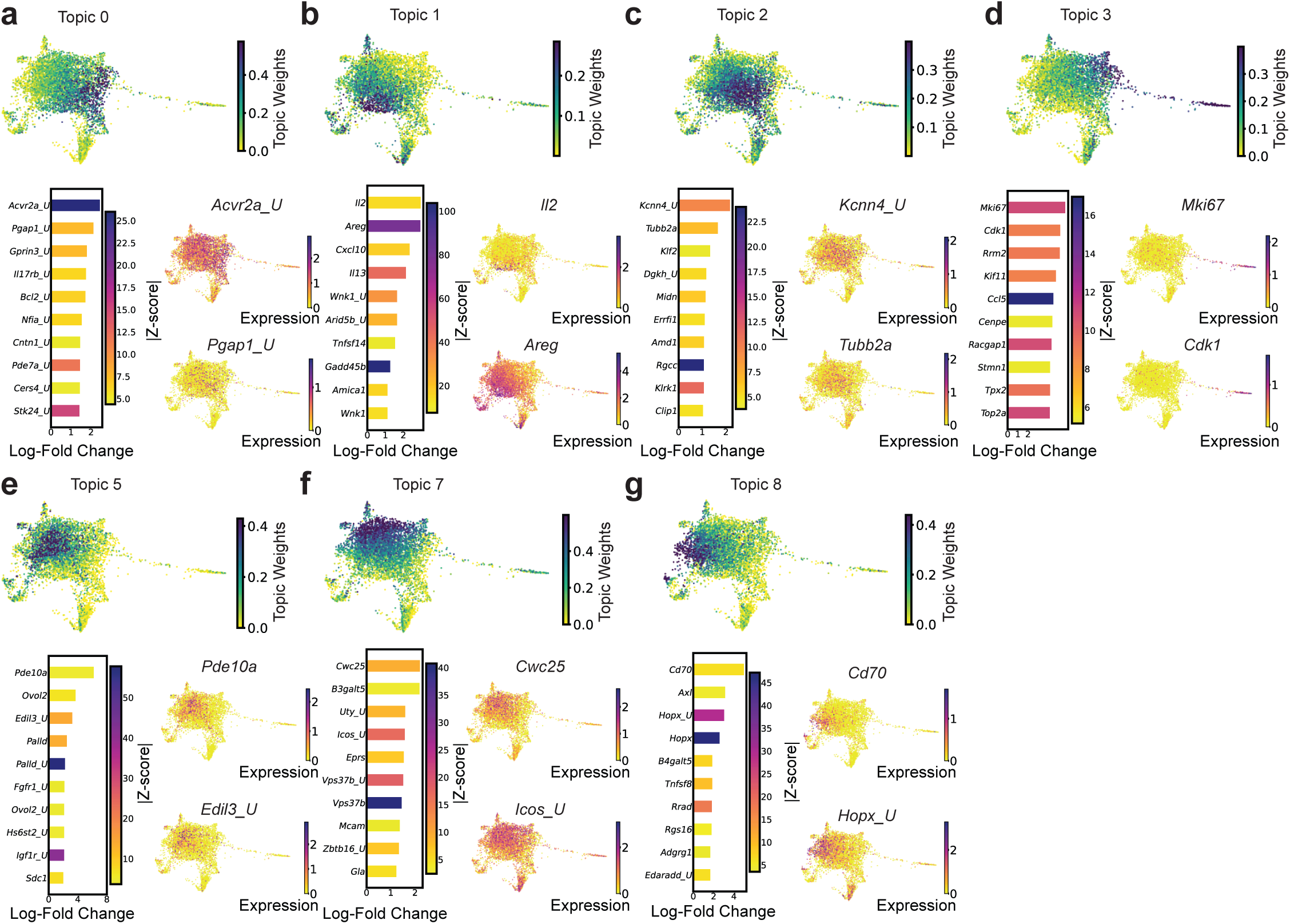
Topic modeling analysis of the ILCs data from only day 3. **a**, For topic 0, force-directed layout (FDL) embeddings shows cells colored by topic weights (top) and by log-normalized expression of topic-specific genes (bottom right); bar plot (bottom left) shows top 10 topic-specific genes ranked by log-fold change (x axis) and colored by absolute value of z-score; *’ U’* indicates unspliced transcripts. **b–g,** Analogous to a, for topics 1–8, respectively.

**Supplementary Figure 8:**
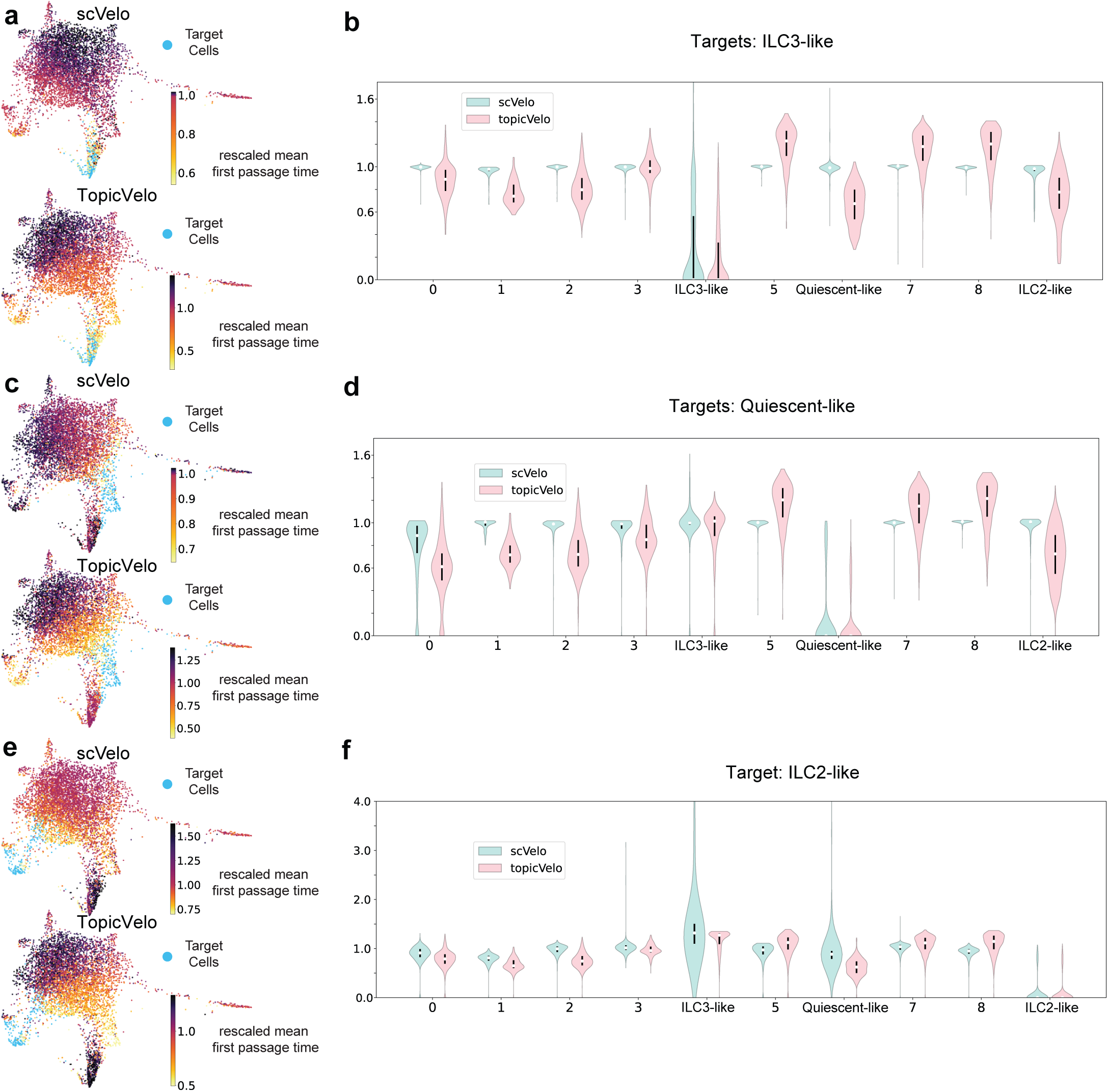
Mean first-passage time analysis of the skin ILCs data. **a, b**, Median-rescaled mean first-passage time (rmfpt) to target group ILC3-like cells. FDL plots (**a**) show cells colored by rmfpt to target group ILC3-like cells (blue), as estimated by *scVelo* (top) and *TopicVelo* (bottom). Violin plots (**b,**) show distributions of rmfpt to ILC3-like cells for subsets of cells, grouped by the topic for which they have highest weight (x axis). **c, d,** Analogous to a, b, with target group set to quiescent-like cells. **e, f,** Analogous to a, b, with target group set to ILC2-like cells.

**Supplementary Figure 9:**
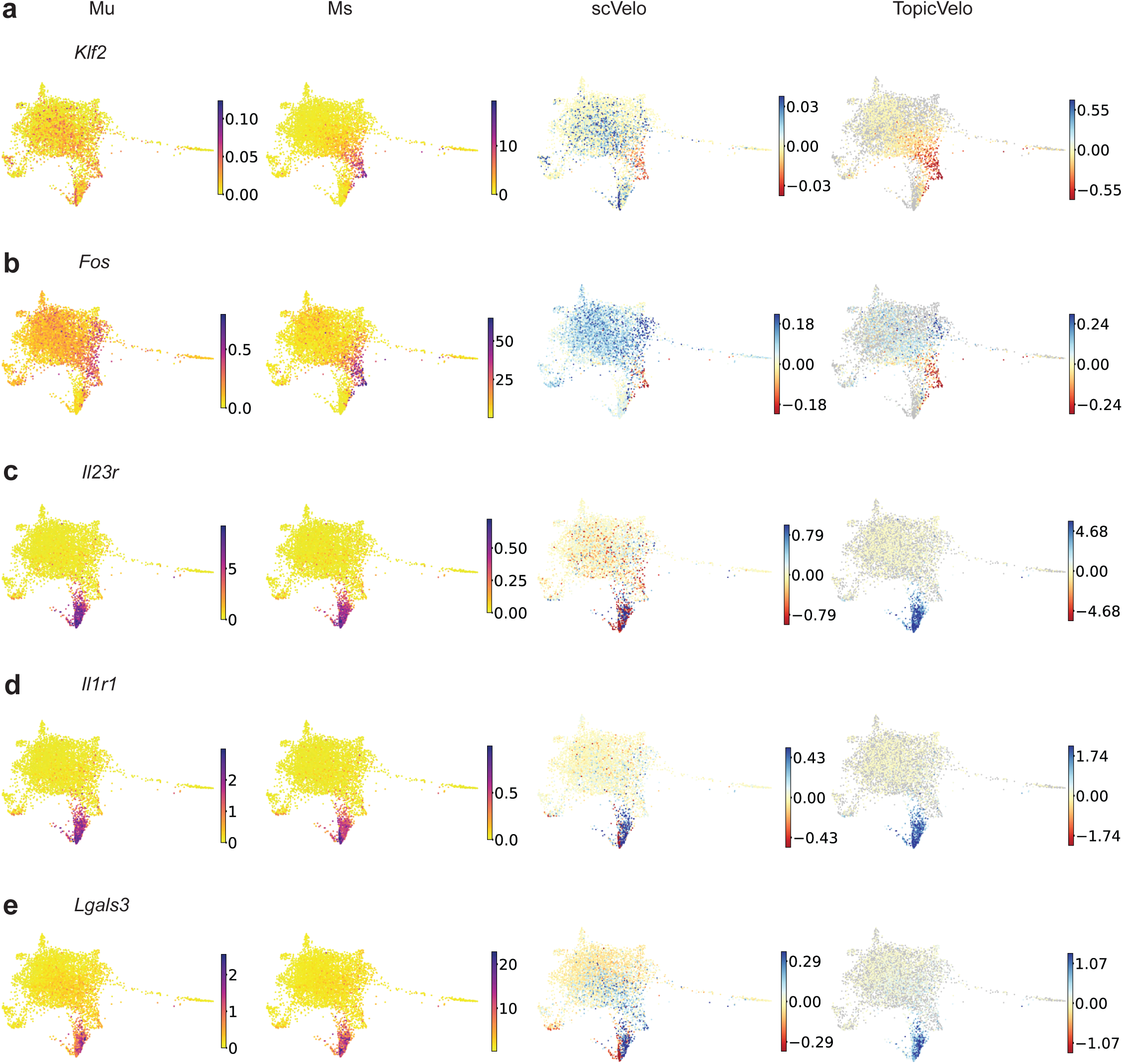
*TopicVelo* recovers more biologically plausible velocities than *scVelo* for the skin ILCs data. **a, b,** Analysis of topic-6 specific genes. FDL plots colored by smoothed size-normalized counts of unspliced (Mu) (far left) and spliced (Ms) (middle left) transcripts, and by velocities inferred by *scVelo* (middle right) and *TopicVelo* (far right), for the genes *Klf2* (**a**) and *Fos* (**b**). **c–e** Analysis of topic-4 specific genes *Il23r*, *Il1r1*, and *Lgals3*, analogous to a, b.

## Notes

### Competing Interest Statement

The authors have declared no competing interest.

## References

1. Kharchenko, P. V. The triumphs and limitations of computational methods for scRNA-seq. Nature Methods 18, 723–732. issn: 1548-7105. https://doi.org/10.1038/s41592-021-01171-x (July 2021).

2. Lähnemann, D., et al. Eleven grand challenges in single-cell data science. Genome Biology 21, 31. issn: 1474-760X. https://doi.org/10.1186/s13059-020-1926-6 (Feb. 2020).

3. Saelens, W., Cannoodt, R., Todorov, H. & Saeys, Y. A comparison of single-cell trajectory inference methods. Nature Biotechnology 37, 547–554. issn: 1546-1696. https://doi.org/10.1038/s41587-019-0071-9 (May 2019).

4. Ginhoux, F., Yalin, A., Dutertre, C. A. & Amit, I. Single-cell immunology: Past, present, and future. Immunity 55, 393–404. issn: 1074-7613. https://www.sciencedirect.com/science/article/pii/S1074761322000863 (2022).

5. Fan, J., Slowikowski, K. & Zhang, F. Single-cell transcriptomics in cancer: computational challenges and opportunities. en. Experimental & Molecular Medicine 52. Number: 9 Publisher: Nature Publishing Group, 1452–1465. issn: 2092-6413. https://www.nature.com/articles/s12276-020-0422-0 (2023) (Sept. 2020).

6. Kunz, D. J., Gomes, T. & James, K. R. Immune Cell Dynamics Unfolded by Single-Cell Technologies. Frontiers in Immunology 9. issn: 1664-3224. https://www.frontiersin.org/articles/10.3389/fimmu.2018.01435 (2023) (2018).

7. La Manno, G. et al. RNA velocity of single cells. Nature 560, 494–498. issn: 1476-4687. https://doi.org/10.1038/s41586-018-0414-6 (Aug. 2018).

8. Bergen, V., Lange, M., Peidli, S., Wolf, F. A. & Theis, F. J. Generalizing RNA velocity to transient cell states through dynamical modeling. Nature Biotechnology, 1546–1696. https://doi.org/10.1038/s41587-020-0591-3 (2020).

9. Wolf, F. A. et al. PAGA: graph abstraction reconciles clustering with trajectory inference through a topology preserving map of single cells. Genome Biology 20, 59. issn: 1474-760X. https://doi.org/10.1186/s13059-019-1663-x (Mar. 2019).

10. Street, K. et al. Slingshot: cell lineage and pseudotime inference for single-cell transcriptomics. BMC Genomics 19, 477. issn: 1471-2164. https://doi.org/10.1186/s12864-018-4772-0 (June 2018).

11. Haghverdi, L., Büttner, M., Wolf, F. A., Buettner, F. & Theis, F. J. Diffusion pseudotime robustly reconstructs lineage branching. Nature Methods 13, 845–848. issn: 1548-7105. https://doi.org/10.1038/nmeth.3971 (Oct. 2016).

12. Barile, M. et al. Coordinated changes in gene expression kinetics underlie both mouse and human erythroid maturation. Genome Biology 22. issn: 1474–760X. https://doi.org/10.1186/s13059-021-02414-y (July 2021).

13. Bergen, V., Soldatov, R. A., Kharchenko, P. V. & Theis, F. J. RNA velocity-current challenges and future perspectives. Molecular Systems Biology 17, e10282. issn: 1744–4292. https://doi.org/10.15252/msb.202110282 (Aug. 2021).

14. Pijuan-Sala, B. et al. A single-cell molecular map of mouse gastrulation and early organogenesis. en. Nature 566. issn: 1476-4687. https://doi.org/10.1038/s41586-019-0933-9(2021) (Feb. 2019).

15. Gorin, G., Fang, M., Chari, T. & Pachter, L. RNA velocity unraveled. PLOS Computational Biology 18, 1–55. https://doi.org/10.1371/journal.pcbi.1010492 (Sept. 2022).

16. Zheng, S. C., Stein-O’Brien, G., Boukas, L., Goff, L. A. & Hansen, K. D. Pumping the brakes on RNA velocity – understanding and interpreting RNA velocity estimates. bioRxiv. eprint: https://www.biorxiv.org/content/early/2022/06/25/2022.06.19.494717.full.pdf. https://www.biorxiv.org/content/early/2022/06/25/2022.06.19.494717 (2022).

17. Gorin, G., Svensson, V. & Pachter, L. Protein velocity and acceleration from single-cell multiomics experiments. Genome Biology 21, 39. issn: 1474-760X. https://doi.org/10.1186/s13059-020-1945-3 (Feb. 2020).

18. Qiu, X. et al. Mapping transcriptomic vector fields of single cells. Cell 185, 690–711.e45. issn: 0092-8674. https://www.sciencedirect.com/science/article/pii/S0092867421015774 (2022).

19. Li, C., Virgilio, M., Collins, K. L. & Welch, J. D. Single-cell multi-omic velocity infers dynamic and decoupled gene regulation. bioRxiv. eprint: https://www.biorxiv.org/content/early/2021/12/15/2021.12.13.472472.full.pdf. https://www.biorxiv.org/content/early/2021/12/15/2021.12.13.472472 (2021).

20. Gorin, G. & Pachter, L. Analysis of Length Biases in Single-Cell RNA Sequencing of Unspliced mRNA by Markov Modeling. Biophysical Journal 120. Publisher: Elsevier, 81a. issn: 0006-3495. https://doi.org/10.1016/j.bpj.2020.11.706 (2021) (Feb. 2021).

21. Lange, M. et al. CellRank for directed single-cell fate mapping. Nature Methods 19, 159–170. https://doi.org/10.1038/s41592-021-01346-6 (2022).

22. Gayoso, A. et al. Deep generative modeling of transcriptional dynamics for RNA velocity analysis in single cells. bioRxiv. eprint: https://www.biorxiv.org/content/early/2022/08/15/2022.08.12.503709.full.pdf. https://www.biorxiv.org/content/early/2022/08/15/2022.08.12.503709 (2022).

23. Qiao, C. & Huang, Y. Representation learning of RNA velocity reveals robust cell transitions. Proceedings of the National Academy of Sciences 118, e2105859118. eprint: https://www.pnas.org/doi/pdf/10.1073/pnas.2105859118. https://www.pnas.org/doi/abs/10.1073/pnas.2105859118 (2021).

24. Gao, M., Qiao, C. & Huang, Y. UniTVelo: temporally unified RNA velocity reinforces single-cell trajectory inference. bioRxiv. eprint: https://www.biorxiv.org/content/early/2022/09/01/2022.04.27.489808.full.pdf. https://www.biorxiv.org/content/early/2022/09/01/2022.04.27.489808 (2022).

25. Farrell, S., Mani, M. & Goyal, S. Inferring single-cell dynamics with structured dynamical representations of RNA velocity. bioRxiv. eprint: https://www.biorxiv.org/content/early/2022/08/23/2022.08.22.504858.full.pdf. https://www.biorxiv.org/content/early/2022/08/23/2022.08.22.504858 (2022).

26. Cui, H., Maan, H., Taylor, M. D. & Wang, B. DeepVelo: Deep Learning extends RNA velocity to multi-lineage systems with cell-specific kinetics. bioRxiv. eprint: https://www.biorxiv.org/content/early/2022/05/30/2022.04.03.486877.full.pdf. https://www.biorxiv.org/content/early/2022/05/30/2022.04.03.486877 (2022).

27. Blei, D. M., Ng, A. Y. & Jordan, M. I. Latent Dirichlet Allocation. Journal of Machine Learning Research 3, 993–1022. (2018) (2003).

28. Blei, D. M. Probabilistic topic models. Science 55, 77–84. https://doi.org/10.1145/2133806.2133826 (2018) (2012).

29. Pritchard, J. K., Stephens, M. & Donnelly, P. Inference of Population Structure Using Multilocus Genotype Data. Genetics 155, 945–959. issn: 1943-2631. https://doi.org/10.1093/genetics/155.2.945(2021) (June 2000).

30. Erosheva, E. A. in Bayesian Statistics 7 (eds Bernardo, J. M., et al.) 501–510 (Oxford University Press, Oxford, 2003).

31. Singh, A. & Bokes, P. Consequences of mRNA Transport on Stochastic Variability in Protein Levels. Biophysical Journal 103, 1087–1096. issn: 0006-3495. https://www.sciencedirect.com/science/article/pii/S0006349512007904 (2012).

32. Setty, M., et al. Characterization of cell fate probabilities in single-cell data with Palantir. Nature Biotechnology 37. Publisher: Nature Publishing Group, 451–460. issn: 15461696. https://doi.org/10.1038/s41587-019-0068-4 (Apr. 2019).

33. Bielecki, P. et al. Skin-resident innate lymphoid cells converge on a pathogenic effector state. Nature 592, 128–132. issn: 1476-4687. https://doi.org/10.1038/s41586-021-03188-w (Apr. 2021).

34. Levens, D. & Larson, D. R. A new twist on transcriptional bursting. eng. Cell 158. S0092–8674(14)00869-1[PII], 241–242. issn: 1097-4172. https://doi.org/10.1016/j.cell.2014.06.042 (July 2014).

35. Dey, K. K., Hsiao, C. J. & Stephens, M. Visualizing the structure of RNA-seq expression data using grade of membership models. PLoS genetics 13, e1006599. issn: 1553-7404. https://doi.org/10.1371/journal.pgen.1006599 (Mar. 2017).

36. Zhao, Y., Cai, H., Zhang, Z., Tang, J. & Li, Y. Learning interpretable cellular and gene signature embeddings from single-cell transcriptomic data. Nature Communications 12, 5261. issn: 2041-1723. https://doi.org/10.1038/s41467-021-25534-2 (Sept. 2021).

37. Carbonetto, P. et al. Interpreting structure in sequence count data with differential expression analysis allowing for grades of membership. bioRxiv. eprint: https://www.biorxiv.org/content/early/2023/03/06/2023.03.03.531029.full.pdf. https://www.biorxiv.org/content/early/2023/03/06/2023.03.03.531029 (2023).

38. Carbonetto, P., Sarkar, A., Wang, Z. & Stephens, M. Non-negative matrix factorization algorithms greatly improve topic model fits. arXiv 2105.13440. arXiv: 2105.13440. https://arxiv.org/abs/2105.13440 (2021).

39. Cao, J., Xia, T., Li, J., Zhang, Y. & Tang, S. A density-based method for adaptive LDA model selection. *Neurocomputing* **72**. Advances in Machine Learning and Computational Intelligence, 1775–1781. issn: 0925-2312. https://www.sciencedirect.com/science/article/pii/S092523120800372X (2009).

40. Deveaud, R., SanJuan, E. & Bellot, P. Accurate and effective latent concept modeling for ad hoc information retrieval. Document numérique 17, 61–84 (Apr. 2014).

41. Röder, M., Both, A. & Hinneburg, A. *Exploring the Space of Topic Coherence Measures* in *Proceedings of the Eighth ACM International Conference on Web Search and Data Mining* (Association for Computing Machinery, Shanghai, China, 2015), 399–408. isbn: 9781450333177. https://doi.org/10.1145/2684822.2685324.

42. Gillespie, D. T. A general method for numerically simulating the stochastic time evolution of coupled chemical reactions. Journal of Computational Physics 22, 403–434. issn: 0021-9991. https://www.sciencedirect.com/science/article/pii/0021999176900413 (1976).

43. Virtanen, P. et al. SciPy 1.0: Fundamental Algorithms for Scientific Computing in Python. Nature Methods 17, 261–272 (2020).

44. Erhard, F. et al. Time-resolved single-cell RNA-seq using metabolic RNA labelling. Nature Reviews Methods Primers 2, 77. issn: 2662-8449. https://doi.org/10.1038/s43586-022-00157-z (Sept. 2022).

45. Songdej, N. et al. Transcription Factor RUNX1 Regulates Factor FXIIIA Subunit (F13A1) Expression in Megakaryocytic Cells and Platelet F13A1 Expression is Downregulated in RUNX1 Haplodeficiency. Blood 136, 25–26. issn: 0006-4971. https://doi.org/10.1182/blood-2020-141382 (Nov. 2020).

46. Psaila, B. et al. Single-Cell Analyses Reveal Megakaryocyte-Biased Hematopoiesis in Myelofibrosis and Identify Mutant Clone-Specific Targets. Molecular Cell 78, 477–492.e8. issn: 1097-2765. https://www.sciencedirect.com/science/article/pii/S1097276520302343 (2020).

47. Shim, M.-H., Hoover, A., Blake, N., Drachman, J. G. & Reems, J. A. Gene expression profile of primary human CD34+CD38lo cells differentiating along the megakaryocyte lineage. Experimental Hematology 32, 638–648. https://doi.org/10.1016/j.exphem.2004.04.002 (July 2004).

48. Li, Y., Qi, X., Liu, B. & Huang, H. The STAT5-GATA2 pathway is critical in basophil and mast cell differentiation and maintenance. en. J Immunol 194, 4328–4338 (Mar. 2015).

49. Karlsson, M. et al. A single-cell type transcriptomics map of human tissues. Science Advances 7. https://doi.org/10.1126/sciadv.abh2169 (July 2021).

50. Cvejic, A. et al. SMIM1 underlies the Vel blood group and influences red blood cell traits. Nature Genetics 45, 542–545. issn: 1546–1718. https://doi.org/10.1038/ng.2603 (May 2013).

51. Suzuki, M. et al. GATA factor switching from GATA2 to GATA1 contributes to erythroid differentiation. Genes to Cells 18, 921–933. eprint: https://onlinelibrary.wiley.com/doi/pdf/10.1111/gtc.12086. https://onlinelibrary.wiley.com/doi/abs/10.1111/gtc.12086 (2013).

52. Tusi, B. K. et al. Population snapshots predict early haematopoietic and erythroid hierarchies. en. Nature 555, 54–60 (Feb. 2018).

53. Yan, H. et al. Developmental differences between neonatal and adult human erythropoiesis. American Journal of Hematology 93, 494–503. eprint: https://onlinelibrary.wiley.com/doi/pdf/10.1002/ajh.25015. https://onlinelibrary.wiley.com/doi/abs/10.1002/ajh.25015 (2018).

54. Sichien, D. et al. IRF8 Transcription Factor Controls Survival and Function of Terminally Differentiated Conventional and Plasmacytoid Dendritic Cells, Respectively. en. Immunity 45, 626–640 (Sept. 2016).

55. Wang, H. et al. Decoding Human Megakaryocyte Development. Cell Stem Cell 28, 535–549.e8. issn: 1934-5909. https://www.sciencedirect.com/science/article/pii/S1934590920305440 (2021).

56. Pellin, D. et al. A comprehensive single cell transcriptional landscape of human hematopoietic progenitors. Nature Communications 10, 2395. issn: 2041-1723. https://doi.org/10.1038/s41467-019-10291-0 (June 2019).

57. Chertov, O. et al. Identification of human neutrophil-derived cathepsin G and azurocidin/CAP37 as chemoattractants for mononuclear cells and neutrophils. en. J Exp Med 186, 739–747 (Aug. 1997).

58. Colonna, M. Innate Lymphoid Cells: Diversity, Plasticity, and Unique Functions in Immunity. Immunity 48, 1104–1117. https://doi.org/10.1016/j.immuni.2018.05.013 (2018).

59. O’Shea, J. J. & Paul, W. E. Mechanisms Underlying Lineage Commitment and Plasticity of Helper CD4¡sup¿+¡/sup¿ T Cells. Science 327, 1098–1102. eprint: https://www.science.org/doi/pdf/10.1126/science.1178334. https://www.science.org/doi/abs/10.1126/science.1178334 (2010).

60. Vivier, E. et al. Innate Lymphoid Cells: 10 Years On. en. Cell 174, 1054–1066 (Aug. 2018).

61. Cao, Z., Sun, X., Icli, B., Wara, A. K. & Feinberg, M. W. Role of Kruppel-like factors in leukocyte development, function, and disease. en. Blood 116, 4404–4414 (July 2010).

62. Gu, Y., Blaauw, D. & Welch, J. D. Bayesian Inference of RNA Velocity from Multi-Lineage Single-Cell Data. bioRxiv. eprint: https://www.biorxiv.org/content/early/2022/07/10/2022.07.08.499381.full.pdf. https://www.biorxiv.org/content/early/2022/07/10/2022.07.08.499381 (2022).

63. Qin, Q., Bingham, E., Manno, G. L., Langenau, D. M. & Pinello, L. Pyro-Velocity: Probabilistic RNA Velocity inference from single-cell data. bioRxiv. eprint: https://www.biorxiv.org/content/early/2022/10/14/2022.09.12.507691.full.pdf. https://www.biorxiv.org/content/early/2022/10/14/2022.09.12.507691 (2022).

64. Gorin, G. & Pachter, L. Length Biases in Single-Cell RNA Sequencing of pre-mRNA. bioRxiv, 2021.07.30.454514. https://doi.org/10.1101/2021.07.30.454514 (July 2021).

65. Lambert, S. A. et al. The Human Transcription Factors. Cell 172, 650–665. issn: 0092-8674. https://www.sciencedirect.com/science/article/pii/S0092867418301065 (2018).

66. Pokhilko, A. et al. Targeted single-cell RNA sequencing of transcription factors enhances the identification of cell types and trajectories. en. Genome Res. 31, 1069–1081 (June 2021).

67. Vayansky, I. & Kumar, S. A. A review of topic modeling methods. Information Systems 94, 101582. issn: 0306-4379. https://www.sciencedirect.com/science/article/pii/S0306437920300703 (2020).

68. Gorin, G., Vastola, J. J., Fang, M. & Pachter, L. Interpretable and tractable models of transcriptional noise for the rational design of single-molecule quantification experiments. bioRxiv. eprint: https://www.biorxiv.org/content/early/2021/12/26/2021.09.06.459173.full.pdf. https://www.biorxiv.org/content/early/2021/12/26/2021.09.06.459173 (2021).

69. Soneson, C., Srivastava, A., Patro, R. & Stadler, M. B. Preprocessing choices affect RNA velocity results for droplet scRNA-seq data. PLOS Computational Biology 17, 1–26. https://doi.org/10.1371/journal.pcbi.1008585 (Jan. 2021).

70. Chen, Z., King, W. C., Hwang, A., Gerstein, M. & Zhang, J. ¡i¿DeepVelo¡/i¿: Single-cell transcriptomic deep velocity field learning with neural ordinary differential equations. Science Advances 8, eabq3745. eprint: https://www.science.org/doi/pdf/10.1126/sciadv.abq3745. https://www.science.org/doi/abs/10.1126/sciadv.abq3745 (2022).

71. Lee, M. *bab2min/tomotopy: 0.12.3* version v0.12.3. July 2022. https://doi.org/10.5281/zenodo.6868418.

72. Lee, D. D. & Seung, H. S. Learning the parts of objects by non-negative matrix factorization. Nature 401, 788–791. issn: 1476-4687. https://doi.org/10.1038/44565 (Oct. 1999).

73. Stephens, M. False discovery rates: a new deal. Biostatistics 18, 275–294. issn: 1465-4644. eprint: https://academic.oup.com/biostatistics/article-pdf/18/2/275/11057424/kxw041.pdf. https://doi.org/10.1093/biostatistics/kxw041 (Oct. 2016).

74. Li, T., Shi, J., Wu, Y. & Zhou, P. On the Mathematics of RNA Velocity I: Theoretical Analysis. CSIAM Transactions on Applied Mathematics 2, 1–55. issn: 2708-0579. http://global-sci.org/intro/article_detail/csiam-am/18653.html (2021).

75. Lam, S. K., Pitrou, A. & Seibert, S. Numba: A llvm-based python jit compiler in Proceedings of the Second Workshop on the LLVM Compiler Infrastructure in HPC (2015), 1–6.

76. Hochgerner, H., Zeisel, A., Lönnerberg, P. & Linnarsson, S. Conserved properties of dentate gyrus neurogenesis across postnatal development revealed by single-cell RNA sequencing. Nature Neuroscience 21, 290–299. issn: 1546-1726. https://doi.org/10.1038/s41593-017-0056-2 (Feb. 2018).

